# Astrocyte transcriptomic analysis identifies glypican 5 downregulation as a contributor to synaptic dysfunction in Alzheimer’s disease models

**DOI:** 10.1101/2024.10.30.621182

**Authors:** Isabel H. Salas, Adrien Paumier, Tao Tao, Aksinya Derevyanko, Corey Switzler, Jillybeth Burgado, Melia Movsesian, Setareh Metanat, Trinity Dawoodtabar, Quinn Asbell, Arash Fassihi, Nicola J. Allen

**Affiliations:** Salk Institute for Biological Studies, Molecular Neurobiology Laboratory, 10010 North Torrey Pines Road, La Jolla, CA 92037, USA; University of California San Diego, 9500 Gilman Drive, La Jolla, CA 92093, USA

**Keywords:** Astrocytes, Glia, Alzheimer’s disease, Neurodegenerative disorders, Synapses, Learning and memory, Synaptic transmission, Dementia, Cognitive decline

## Abstract

Synaptic dysfunction is an early feature in Alzheimer’s disease (AD) and correlates with cognitive decline. Astrocytes are essential regulators of synapses, impacting synapse formation, maturation, elimination and function. To understand if synapse-supportive functions of astrocytes are altered in AD, we used astrocyte BacTRAP mice to generate a comprehensive dataset of hippocampal astrocyte transcriptional alterations in two mouse models of Alzheimer’s pathology (APPswe/PS1dE9 and Tau P301S), characterizing sex and age-dependent changes. We found that astrocytes from both models downregulate genes important for synapse regulation and function such as the synapse-maturation factor Glypican 5. This transcriptional signature is shared with human post-mortem AD patients. Manipulating a key component of this signature by *in vivo* overexpression of Glypican 5 in astrocytes is sufficient to prevent early synaptic dysfunction and improve spatial learning in APPswe/PS1dE9 mice. These findings open new avenues to target astrocytic factors to mitigate AD synaptic dysfunction.

## INTRODUCTION

Alzheimer’s disease (AD) is a neurodegenerative disorder characterized by progressive neuronal loss and decline of cognitive function^1^. Pathologically, AD brains present misfolded protein aggregations including extracellular amyloid-beta plaques and intracellular neurofibrillary tangles, composed of hyperphosphorylated tau^2,3^.These aggregates spread progressively in the brain with hippocampus and prefrontal cortex being the regions first and most affected^4^. Another major pathological feature of AD is synapse dysfunction and loss, which is the strongest correlate of cognitive decline^5,6^. In the hippocampus, synaptic dysfunction follows a progressive pattern characterized by an early neuronal hyperactivity which occurs before dementia onset, followed by a synaptic hypoactivity and neuronal loss^7–9^. This progressive synaptic dysfunction is recapitulated in AD mouse models^10,11^. Research to study synapse dysfunction in AD has long focused on neurons. However, it has become increasingly clear that non-neuronal glial cells, including astrocytes, microglia and oligodendrocytes, play crucial roles in regulating synapse function and might therefore be fundamental in the progression of dementia^12–14^.

Astrocytes are a type of glial cell that play an integral role in the regulation of synaptic function^15^. Astrocytes interact with synapses via processes that contact pre and postsynaptic terminals forming part of the tripartite synapse^16^. They provide nutritional and structural support to synapses, and maintain neuronal excitability by regulating ion homeostasis and neurotransmitter recycling^17,18^. Further, astrocytes produce proteins including glypicans 4 and 6 and thrombospondins that promote the formation of new synapses^19,20^, as well as chordin like 1 and glypican 5 that induce the maturation of existing synapses^21,22^. Additionally, astrocytes participate in synapse elimination through phagocytosis via Megf10, Mertk and the complement pathway^23,24^. In the healthy brain, the regulation of synapse formation, maturation and elimination by astrocytes is essential for synaptic plasticity and for maintenance of learning and memory capabilities, properties that are altered in AD^25^. Thus, understanding how astrocyte synapse-regulating functions are altered in AD brains, and how these changes contribute to synaptic dysfunction and cognitive decline, may open new therapeutic targets to ameliorate dementia.

In AD, astrocytes undergo morphological and transcriptional changes that reflect a reactive state, including upregulation of the cytoskeletal protein GFAP^26,27^. It is now appreciated that astrocyte reactivity comprises a number of different states that are proposed to be protective or detrimental, possibly depending on disease stage^13,14,26,27^. However, how astrocyte alterations in AD affect synaptic function is still poorly understood. Transcriptional studies and *in vitro* experiments suggest a detrimental role of reactive astrocytes in AD, with reduced synaptic support, however *in vivo* astrocyte-specific functional studies are limited^12,13,28^. A further critical question is to understand how astrocyte alterations evolve over time in the progression of AD, as synaptic alterations differ depending on disease stage^7,14^.

In this manuscript we performed an in-depth characterization of age and sex-dependent astrocyte transcriptional changes in mouse models presenting two main pathological hallmarks of AD, amyloidosis and tau pathology. We achieved this by using the translating ribosomal affinity purification (TRAP) approach to tag astrocyte ribosomes in APPswe/PS1dE9 and Tau P301S at three different disease stages and in both sexes (Figures 1-5). In the second part of the manuscript (Figures 6-7) we investigated the functional consequences of astrocyte transcriptional alterations for AD synaptic dysfunction, by genetically manipulating the expression of the synapse-maturation factor Glypican 5 (*Gpc5*), which is downregulated in AD patients and mouse models. We found that *in vivo* overexpression of GPC5 in astrocytes was able to prevent early synaptic hippocampal hyperactivity, as well as improve spatial learning, in the APPswe/PS1dE9 mouse model.

**Figure 1:**
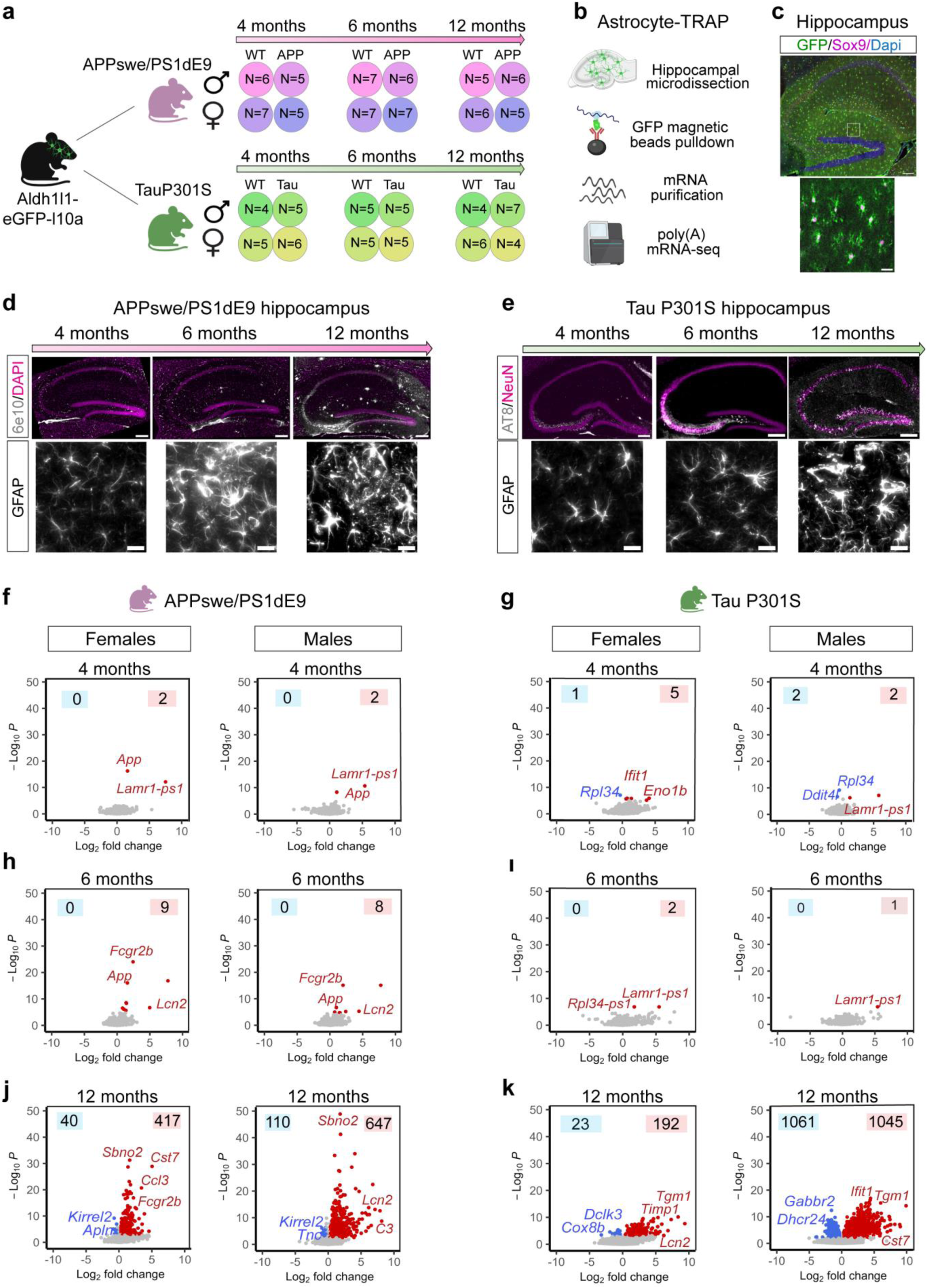
Astrocyte transcriptome is altered in advanced pathology stages in AD mouse models. a) Experimental design for the astrocyte mRNA enrichment approach by crossing the astrocyte BAC-TRAP genetic mouse model Aldh1l1-eGFP-L10a with AD mouse models APPswePS1/dE9 and Tau P301S. N represents the number of mice per group, with a total of 133 mouse hippocampal samples. b) Explanatory diagram for the astrocyte BAC-TRAP ribosomal pulldown followed by mRNA purification and sequencing. c) Representative image of hippocampal astrocytes marked with Sox9 expressing the GFP tag in Aldh1l1-eGFP-L10a mice. Scale bar = 100 µm top panel, 20 µm bottom zoom-in panel. d) Representative images of progressive pathology in *APPswe/PS1dE9* at different ages. Top panels show amyloid plaque accumulation stained with 6e10 antibody (scale bar = 200 µm). Botton panels illustrate increased astrocyte reactivity by GFAP stainings (same GFAP images are shown in Supplementary Fig 1). Scale bar= 20 µm. e) Representative images of progressive pathology in *TauP301S* mice at different ages. Top panels show pathological Tau phosphorylation levels stained with AT8 antibody (scale bar = 200 µm). Bottom panels illustrate increased astrocyte reactivity shown by GFAP staining (same GFAP images are also shown in Supplementary Fig 1). Scale bar= 20 µm. f-k) Volcano plots showing number of differentially expressed genes in different ages, sex and disease models (APP left, Tau right). Top left and right corners show the number of down or upregulated genes for each age and group. Upregulated genes shown in red, downregulated genes in blue. Differentially expressed genes cut-off: *padj* < 0.05, Mean transcripts per million (TPM) > 1, Log2 Fold change (Log2FC) > 0.3(upregulation) or < -0.3 (downregulation). Complete bulk mRNA sequencing data for total of 72 samples for APP and 61 samples for Tau is summarized in Supplementary Tables 2-5.

## RESULTS

### 1. Astrocyte transcriptome is altered in advanced pathology stages in AD mouse models

Our goal is to determine how astrocytes are impacted by two different drivers of AD pathology, amyloid and tau, across disease progression, and in both males and females. To address this we used the APPswe/PS1dE9 mouse line (referred to as APP)^29^, and the Tau P301S line (referred to as Tau)^30^ (**Figure 1a**) as a model of amyloidosis and tau pathology respectively, focusing on the hippocampus, a brain region highly affected in AD. APP mice display amyloid plaques in the hippocampus at 6 months of age, which are widespread by 12 months and accompanied by increased GFAP immunoreactivity, reflecting reactive astrogliosis, starting at 6 months (**Fig 1d, Supplementary Fig 1a-c**). Western blot analysis revealed a significant increase in expression of synaptic proteins at 4 months in APP male mice, with a later downregulation of synaptic proteins at 6 and 12 months in both sexes (**Supplementary Fig 1g**). In the Tau model we observed a mild increase in AT8 immunoreactivity at 6 months, reflecting pathological tau hyperphosphorylation, which is robustly spread by 12 months. Reactive astrogliosis was only observed in females at 12 months (**Supplementary Fig 1d-f**). In contrast, significant synaptic alterations were only present in Tau male mice (**Supplementary Fig 1h**). Based on these findings we selected the following timepoints to determine how the hippocampal astrocyte transcriptome is impacted at three crucial stages of pathology progression: 4 months, before misfolded protein aggregation is present; 6 months, when amyloid plaques and hyperphosphorylated tau deposits begin to appear; 12 months, when pathology is widespread. We profiled astrocytes from male and female mice independently to assess for potential sex differences in the astrocyte transcriptome in AD models (**Fig 1a**).

To enrich for actively translated astrocyte mRNA, we employed the TRAP technique. We crossed the *Aldh1l1-eGFP-Rpl10a* transgenic mouse line, which expresses a GFP-tagged ribosomal protein (L10) under the astrocyte-specific Aldh1l1 promoter^31^, with APP and Tau lines (**Fig 1a**). This enables affinity purification of GFP-tagged ribosomes and associated mRNA followed by mRNA extraction and RNA sequencing (**Fig 1b**). Using immunohistochemistry, we confirmed the efficiency and specificity of the Aldh1l1-eGFP-Rpl10a line, with more than 95% of astrocytes in the hippocampus expressing GFP, and only 3.4% of non-astrocyte cells showing expression (**Supplementary Fig 2a-c**). We performed bulk RNA sequencing from a total of 133 samples encompassing three timepoints, two genotypes and associated wildtype (WT) controls, and both sexes (**Fig 1a**). We confirmed an enrichment of astrocyte-specific genes, along with a depletion of other cell marker genes in the astrocyte pulldown fraction relative to the input (before the pulldown) (**Supplementary Fig 2d,e**). Differential gene expression analysis comparing transgenic and WT mice revealed that in early stages of pathology (4 and 6 months), astrocytes display very few differentially expressed genes (DEGs) in both APP and Tau mouse models (**Fig 1 f-i, Supplementary Tables 2-5**). In contrast, at 12 months of age, both APP and Tau model astrocytes show hundreds of DEGs, with male mice showing a greater number of DEGs compared to females (**Fig 1j,k**). In summary, we have generated a dataset of transcriptional changes of astrocytes across AD pathology and identified robust transcriptional alterations in advanced pathology stages in both amyloid and Tau models.

### 2. Sex differences in astrocyte transcriptional alterations in amyloid model mice

We asked whether the astrocyte gene expression changes in AD models are different between male and female mice, focusing on the 12-month timepoint when both APP and Tau show the highest number of DEGs (**Fig 1**). In the APP model, we observed 245 genes commonly changed in both sexes, while 512 and 212 DEGs were exclusively changed in male and female mice respectively. (**Fig 2a, Supplementary Table 6**). We determined if the direction of DEGs changes are conserved between male and female mice, independently of their significance, by running a Log2 fold change correlation analysis of all expressed genes. We found that the male-female correlation, albeit significant, was not very strong (Spearman rho=0.27, *p*=2.2e-16). In addition, we identified specific genes that are exclusively changed in only one sex (**Fig 2a,c**). To investigate the function of the significantly altered genes we ran over representation analysis with DEGs common to male and female mice, or only altered in one sex. We found an upregulation of immune-related pathways for commonly changed genes, a male-specific upregulation of genes involved in antiviral response, and a female-specific upregulation of genes involved in collagen trimerization and biosynthesis (**Fig 2e, Supplementary Table 6**). Sex-genotype interaction analysis revealed 20 significant genes, 18 of them showing an exacerbated upregulation in male compared to female mice, and 13 of these genes being involved in the immune response (**Fig 2g, Supplementary Table 7**).

**Figure 2:**
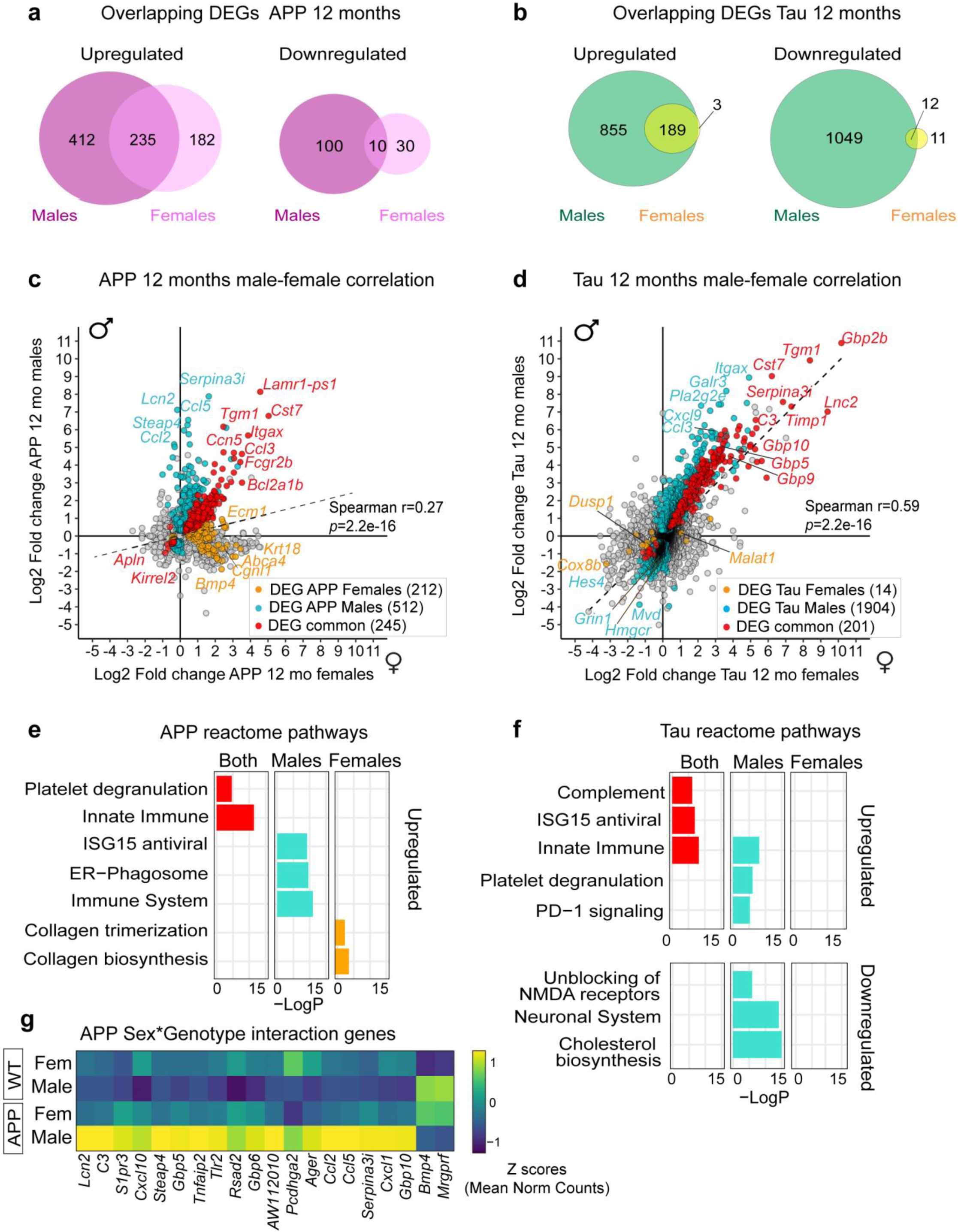
Sex differences in astrocyte transcriptional alterations in amyloid model mice. a-b) Venn diagram showing significant overlapping upregulated (left) and downregulated (right) DEGs between male and female in 12-month-old APP (a) or Tau (b) mice. DEGs significance criteria: *padj* < 0.05, TPM > 1, |Log2FC| > 0.3. Complete gene list can be found in Supplementary Table 6. c-d) Correlation between Log2FC of gene expression in 12-month-old males and females in APP(c) and Tau (d) models. Spearman correlation was calculated with all genes expressed (mean TPM>1) and is shown in the right. Blue dots are significant males-specific DEGs, orange dots are significant female-specific DEGs and red dots are genes commonly altered in both sexes. e-f) Over-representation analysis showing reactome pathways enrichment for genes that are differentially upregulated in 12-month-old males (blue), females (orange) or both (red) for APP (e) and Tau (f) models. Affinity propagation package was used to cluster similar reactome pathways for plotting purposes. Complete list of pathways can be found in Supplementary Table 6. g) Heatmap with genes showing a significant sex-genotype interaction in APP 12-month-old group. Colors represent the Z scores from normalized counts. Complete gene list can be found in Supplementary Table 7.

In the Tau model we found that the majority of upregulated DEGs in females are also upregulated in males (98%), whereas only 12 genes are commonly downregulated (**Fig 2b**). Correlation analysis between Tau male and female Log2 fold change values showed a stronger Spearman correlation than the APP group (rho=0.59, *p*=2.2e-16). Although Tau male mice had a greater number of DEGs compared to females (**Fig 1**), almost all the DEGs were changed in the same direction in the females, even if not reaching statistical significance (**Fig 2d**). In addition, no genes showed a significant sex-genotype interaction in the Tau model (**Supplementary Table 7)**. Over representation analysis revealed a common upregulation of immune-related pathways, and a male-specific downregulation of cholesterol biosynthesis and glutamatergic synapse regulation (**Fig 2f**). This data is consistent with our Western blot results showing greater loss of synaptic markers in Tau male compared to female mice (**Supplementary Fig 1)**. In conclusion, we found that astrocytes display mild sex differences in the presence of AD-relevant pathology, mainly in the amyloid model, with astrocytes in male mice showing an exacerbated immune response compared to female mice. In the Tau line, astrocytes in males show a more pronounced downregulation of synapse-related genes with a parallel decrease in synaptic proteins.

### 3. Astrocyte transcriptional alterations occur progressively with AD pathology

Given that only 20 genes in the APP 12-month dataset, and no genes at other timepoints or in the Tau model, showed a significant sex-genotype interaction (**Fig 2g, Supplementary Table 7**), we combined male and female samples for in-depth analysis to ask how astrocytes are impacted with pathology progression in APP and Tau models (**Fig 3a, Supplementary Tables 3-5**). We first explored whether astrocyte DEGs are overlapping or unique in 12-month old APP and Tau, finding 559 and 90 DEGs that are commonly up and downregulated respectively (**Fig 3b, Supplementary Table 5**). We then asked if astrocyte transcriptional alterations that are present at 12 months occur in a progressive manner and are already present at 6 months, despite not being significant. We found that the majority of DEGs present at 12 months in both APP and Tau models (70-86%) are not changed at 6 months (**Fig 3d,g**). For those genes that are altered, we found that 30% and 12% of upregulated DEGs in APP and Tau models respectively, are already upregulated at 6 months (**Fig 3c**). These genes are mainly involved in immune-related responses, suggesting an early astrocyte inflammatory response already at 6 months, particularly in the APP group. Of downregulated DEGs in APP and Tau, 7% and 1% respectively show downregulation at 6 months, and include genes involved in synapse regulation for the APP group, implying an early dysfunction in synapse-regulating responses in APP astrocytes already at 6 months of age (**Fig 3f**). Finally, 4% of downregulated DEGs in the Tau model are initially upregulated at 6 months (**Fig 3h**). This list includes genes involved in the regulation of synaptic transmission which could indicate a potential neuroprotective response at 6 months in the Tau model. Alternatively, this upregulation might reflect an early hippocampal neuronal hyperactivity found in APP and other Tau mouse models^7^. In summary, we find that with pathology progression astrocytes display a gradual upregulation of immune-related pathways and a downregulation of synapse-regulating genes, particularly in the APP model.

**Figure 3:**
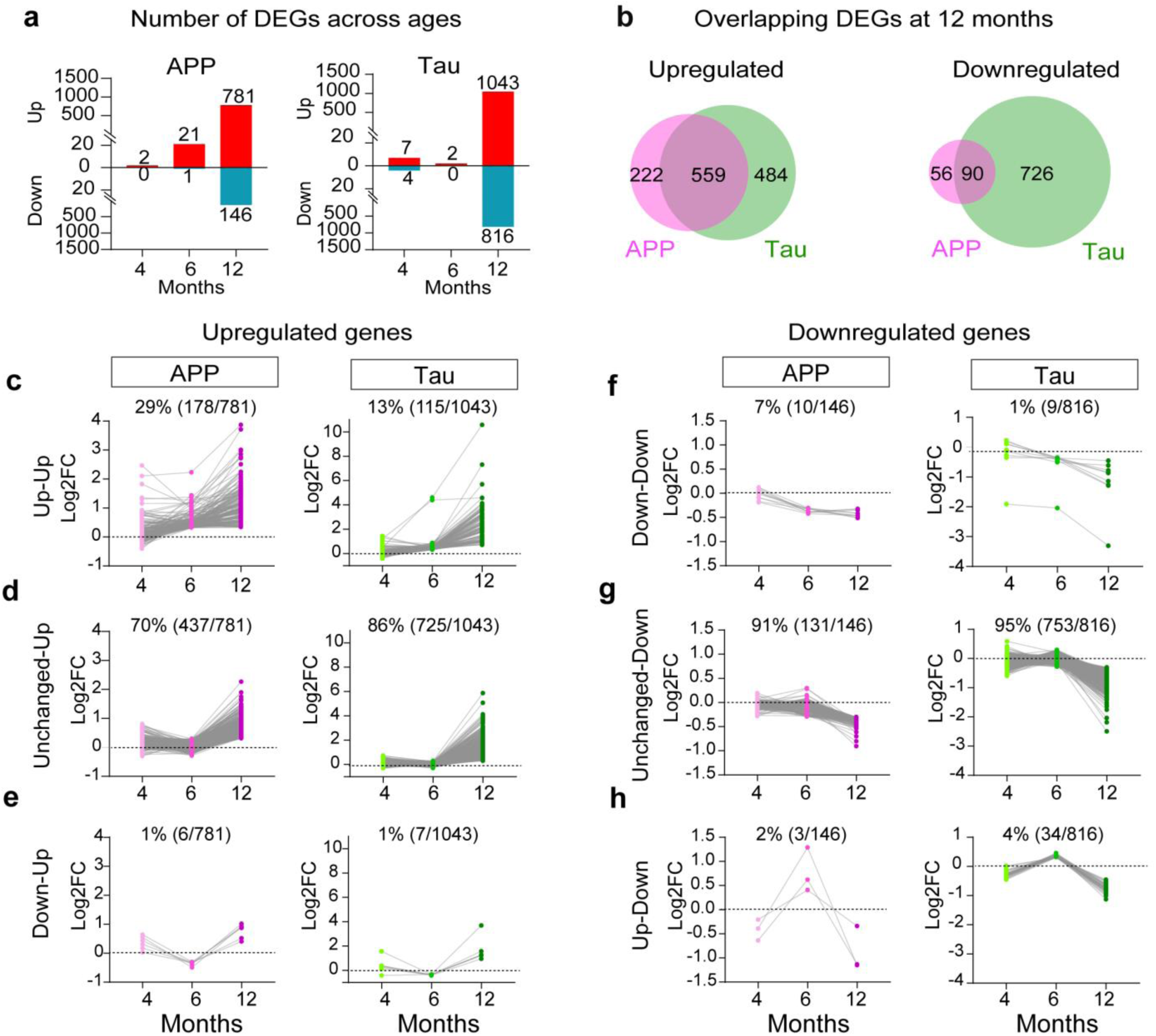
Astrocyte transcriptional alterations occur progressively with AD pathology. a) Number of DEGs combining male and female samples at different ages in APP (left) or Tau (right) mouse models. N=10-14 mice per age, genotype and model (see Fig 1a). DEGs significance criteria: *padj* < 0.05, TPM > 1, |Log2FC| > 0.3. Complete gene list can be found in Supplementary Table 3-5. b) Venn diagram showing overlapping upregulated (left) and downregulated (right) DEGs between 12-month-old APP and Tau groups. c-e) Trajectory of upregulated DEGs at 12 months, categorized by c) Up-up response: early upregulation at 6 months (Log2FC>0.3, independent of significance status); d) Unchanged-up response: genes not changed at 6 months; e) Down-up response: genes downregulated at 6 months (Log2FC<-0.3, independent of significance status). Complete list of genes can be found in Supplementary table 8. f-h) Trajectory of downregulated DEGs at 12 months, categorized by f) Down-down response: early downregulation already at 6 months (Log2FC<-0.3, independent of significance status); g) Unchanged-down response: genes not changed at 6 months; h) Up-Down response: genes upregulated at 6 months (Log2FC>0.3, independent of significance status). Complete list of genes can be found in Supplementary Table 8.

### 4. Genes encoding synapse-related proteins are dysregulated in astrocytes in APP and Tau models

Due to known astrocytic expression of a number of AD risk genes identified in genome-wide association studies (GWAS), we asked if their expression is dysregulated in astrocytes in amyloid or tau pathology models. We first determined the percentage of AD risk genes expressed by astrocytes using three independent GWAS reports^32–34^ (**Fig 4a**, **Supplementary Table 9**). We found that ∼50% of AD risk genes are expressed by astrocytes in our dataset, and ∼10% are enriched in astrocytes (**Fig 4a, Supplementary Fig 3a**). For AD risk genes expressed by astrocytes, we found six genes differentially expressed in 12-month APP and Tau astrocytes. Most of these genes are involved in the regulation of the immune response, including *Bcl3, Trem2, Plcg2, Relb2.* In turn, six genes are changed exclusively in the Tau model, and are mainly related to lipid metabolism such as *Abca1, Abca7* (**Fig 4b, Supplementary Fig 3b-d**). Using single molecule fluorescent *in situ* hybridization we confirmed a 2-fold increase of *Bcl3* mRNA in APP 12-month hippocampal astrocytes and, albeit not significant, a 20% downregulation of *Hbegf* in APP astrocytes compared to WT (**Supplementary Fig 3e-j**).

**Figure 4:**
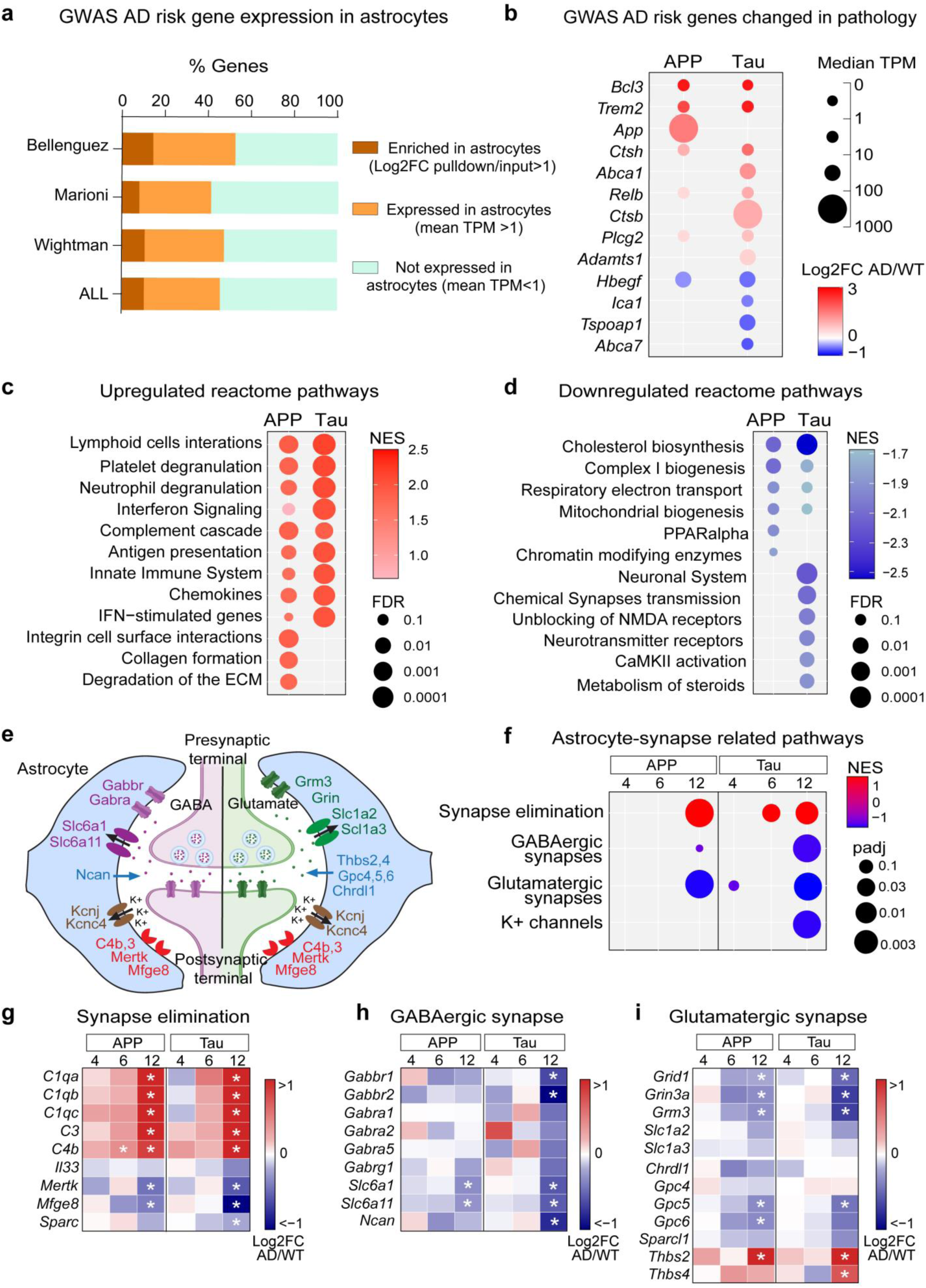
Genes encoding synapse-related proteins are dysregulated in astrocytes in APP and Tau models. a) Percentage of AD risk genes that are either expressed by astrocytes (mean TPM >1), enriched in astrocytes (Log2FC pulldown/input >1; *padj*<0.05), or not expressed in astrocytes (mean TPM<1) for different AD GWAS datasets^32–34^. Complete list of genes can be found in Supplementary Table 9. b) Dot plot showing GWAS AD risk genes that are significantly changed in 12-month-old APP and Tau mice. Colors represent the Log2FC and size of the dots the median TPM values. c-d) Gene set enrichment analysis showing top reactome pathways upregulated (a) and downregulated (b) in astrocytes in 12-month-old APP and Tau mice. Colors represent the normalized enrichment score (NES), size of the dots the significance analyzed by false discovery rate (FDR). Complete list of pathways is included in Supplementary Table 10. e) Diagram showing synapse-related genes in astrocytes. f) Gene set enrichment analysis with custom-made astrocyte synapse-related gene lists for APP and Tau at 4, 6 and 12 months. Colors represent the normalized enrichment score (NES), size of the dots the significance analyzed by *padj* value. Gene sets and complete gene set enrichment analysis is included in Supplementary Table 11. g-i) Heatmaps showing Log2 fold changes of astrocyte genes involved in synapse elimination (g), GABAergic synapse function (h), and glutamatergic synapse function (i). * *padj*<0.05.

**Figure 5:**
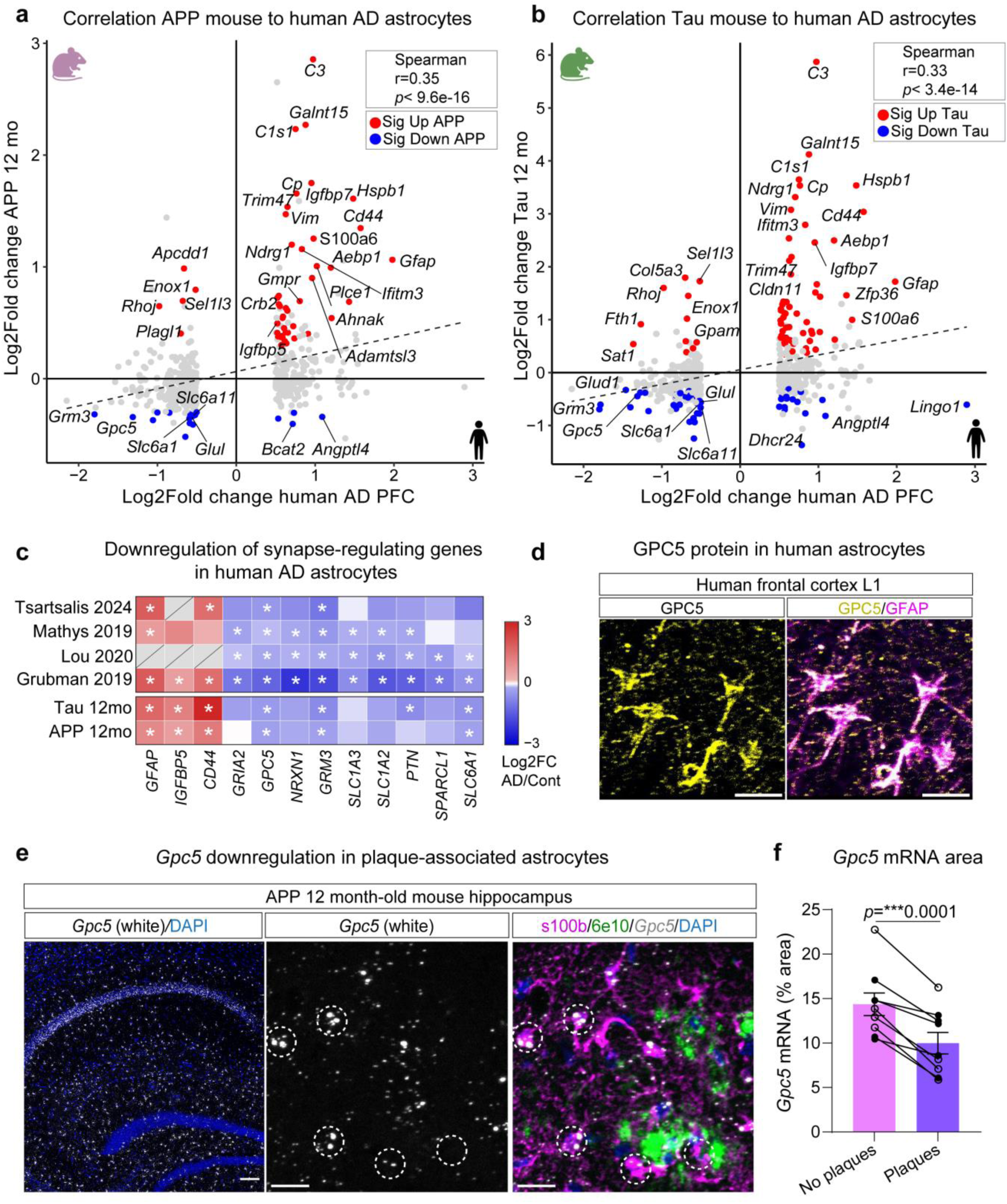
Glypican 5 is downregulated in astrocytes in Alzheimer’s disease patients and mouse models. a-b) Correlation between Log2FC of gene expression in 12-month-old APP (a) and Tau (b) mice and human AD DEGs in astrocytes obtained from Grubman et al.^35^ Red and blue dots depict up and downregulated genes respectively in APP (a) or Tau mice (b). Grey dots are genes exclusively significant in human astrocytes. Complete gene list can be found in Supplementary Table 12. c) Heatmap showing Log2FC in synapse-related genes in astrocytes from postmortem human AD patients and in 12-month-old APP and Tau mice. Human data obtained from^35–38^. * *padj*<0.05. d) Immunostaining of postmortem human frontal cortex showing GPC5 protein expression in astrocytes marked with GFAP in layer 1. Scale bar = 20 µm. e-f) Representative image showing *Gpc5* mRNA *in situ* hybridization in APP 12-month-old hippocampus (left, scale bar = 100 µm). Right: Zoom in panel showing *Gpc5* mRNA levels (white), astrocytes (marked with s100b, magenta) and amyloid plaques (stained with 6e10, green). Scale bar = 20 µm (e). f) Quantification of *Gpc5* mRNA signal (% area) in astrocytes associated with amyloid plaques (plaques) or non-associated with amyloid plaques (no plaques). Each datapoint is a mouse (open circles=female, close=male). *N*= 7 (4F, 3M). Paired t-test was performed for statistical analysis.

Next, to investigate the function of astrocyte transcriptional changes occurring at 12 months of age in APP and Tau models we ran gene set enrichment analysis, ranking genes by Log2 fold change, independent of significance status. We found a strong upregulation of immune-related pathways in both models, and an APP-specific upregulation of extracellular matrix-related pathways (**Fig 4c, Supplementary Table 10**). Both APP and Tau models show a downregulation of genes involved in cholesterol biosynthesis, and Tau mice display a downregulation of pathways involved in synaptic transmission (**Fig 4d**). To further explore whether astrocyte-expressed synapse-regulating factors are altered in AD mouse models, we generated a gene set with previously reported synapse-regulating factors and ran gene-set enrichment analysis (**Fig 4e-f, Supplementary Table 11**). We found that astrocytes in both APP and Tau models show a progressive upregulation in genes associated with synapse elimination, particularly of genes involved in the complement cascade (**Fig 4f,g**). This is accompanied by progressive downregulation of genes that modulate glutamatergic and GABAergic synapses, the latter being more pronounced in the Tau model (**Fig 4f,h**). Downregulated genes include those encoding receptors and transporters (*Slc6a11, Grm3, Grin3a, Grid1*), and factors involved in synapse formation and maturation (*Gpc5, Gpc6, Ncan*) (**Fig 4h,i**). In summary, this data suggest that astrocytes compromise their synapse-related roles in APP and Tau mouse models.

### 5. Glypican 5 is downregulated in astrocytes in Alzheimer’s disease patients and mouse models

As a first step to investigate the functional consequences of astrocyte gene expression changes for synaptic function in AD, we asked whether the transcriptional alterations that we identified in AD mouse models are conserved in human AD patients. For this we compared our results with a human AD single nucleus RNA-sequencing study performed in the entorhinal cortex, which identified 700 DEGs in astrocytes^35^. Of these 700 genes, 550 have mouse orthologs, and 39 and 12 of the genes are commonly up and downregulated respectively in 12-month-old APP and Tau models (**Supplementary Fig 4a, Supplementary Table 12**). Correlation analysis comparing Log2 fold changes of significant DEGs in human AD astrocytes with mouse models, revealed a modest but significant spearman correlation between human and APP astrocytes (*r*=0.35, *p*=9.6e-16) and human and Tau astrocytes (*r*=0.33, *p*=3.4e-14) (**Fig 5a,b**). Among the 12 astrocyte DEGs that are commonly downregulated in human AD patients, APP and Tau mouse models, we found several genes involved in the regulation of synaptic function including *Grm3, Gpc5, Slc6a1, Slc6a11* and *Glud1*. This suggests that astrocytes in both human patients and mouse AD models alter their synapse-supporting functions and may contribute to synaptic deficits in AD. To determine if a downregulation of synapse-regulating genes in astrocytes is observed in additional datasets from human AD patients, we compared our results with three independent human datasets^36–38^, finding a consistent downregulation of genes important for synaptic function (**Fig 5c**). Interestingly, this downregulation of synapse-related genes is also present in reactive astrocyte clusters characterized by a high *GFAP* expression, relative to other clusters. This suggests that the downregulation of synapse-related genes in AD may be triggered by an increase in astrocyte reactivity^39–42^ (**Supplementary Fig 4c**). In fact, this signature is not restricted to AD and occurs in other neurodegenerative disorders including multiple sclerosis (MS) and frontotemporal dementia (FTD)^43,44^ (**Supplementary Fig 4b**).

One of the genes that is significantly downregulated in APP and Tau models, and in human AD, MS and FTD patients is the astrocyte-secreted factor Glypican 5 (*GPC5*)^35–38,43,44^. GPC5 has been previously identified as a regulator of synapse maturation and stabilization in the adult brain, by modulating presynaptic terminal size and postsynaptic recruitment of GluA2-containing AMPARs^22^. Hence, we hypothesized that GPC5 downregulation may contribute to synaptic dysfunction in diverse neurodegenerative disorders. Using immunohistochemistry in human postmortem tissue, we confirmed presence of GPC5 protein in human cortical astrocytes (**Fig 5d, Supplementary Fig 4d**). We then validated the human findings in AD mouse models, focusing on the APP model as it shows an early and consistent increase in GFAP around amyloid plaques (**Supplementary Fig 1**), and a progressive *Gpc5* downregulation when comparing RNA sequencing from 4, 6 and 12 months (**Fig 4i**, **Supplementary Fig 4e**). We performed single molecule RNA *in situ* hybridization for *Gpc5* combined with immunofluorescence to stain for amyloid plaques and the astrocyte marker s100b in APP 12-month mice (**Fig 5e**). We found that *Gpc5* expression is significantly downregulated in Gfap-high reactive astrocytes around amyloid plaques compared to astrocytes not associated with amyloid plaques (**Fig 5f**). These data support a role for astrocyte *Gpc5* downregulation in synaptic dysfunction in neurodegeneration.

### 6. Astrocyte Glypican 5 overexpression prevents hippocampal synaptic hyperactivity in APP mice at 4 months

Astrocyte GPC5 regulates synapse maturation and stabilization, and its expression is downregulated in AD brains. Thus, we hypothesized that astrocyte *Gpc5* downregulation contributes to synaptic dysfunction in neurodegenerative disorders by altering synaptic function. To test this, we prevented the decrease in *Gpc5* by virally overexpressing GPC5 in astrocytes *in vivo* in APP mice, and assayed key features of synaptic transmission and learning and memory that are altered in this model. GPC5 was delivered by retro-orbitally injecting 2-month-old mice with AAV-PHP.eB-HA-Gpc5 (GPC5) or AAV-PHP.eB-smFP-HA as a control, both driven by the minimal GfaABC1D promoter to restrict expression to astrocytes, allowing for brain wide delivery (**Fig 6a, Supplementary Fig 6**). We confirmed that viral overexpression from 2 months to 4 months of age targeted ∼70% of astrocytes in the hippocampus without affecting GFAP immunoreactivity (**Supplementary Fig 5a-c**).

**Figure 6:**
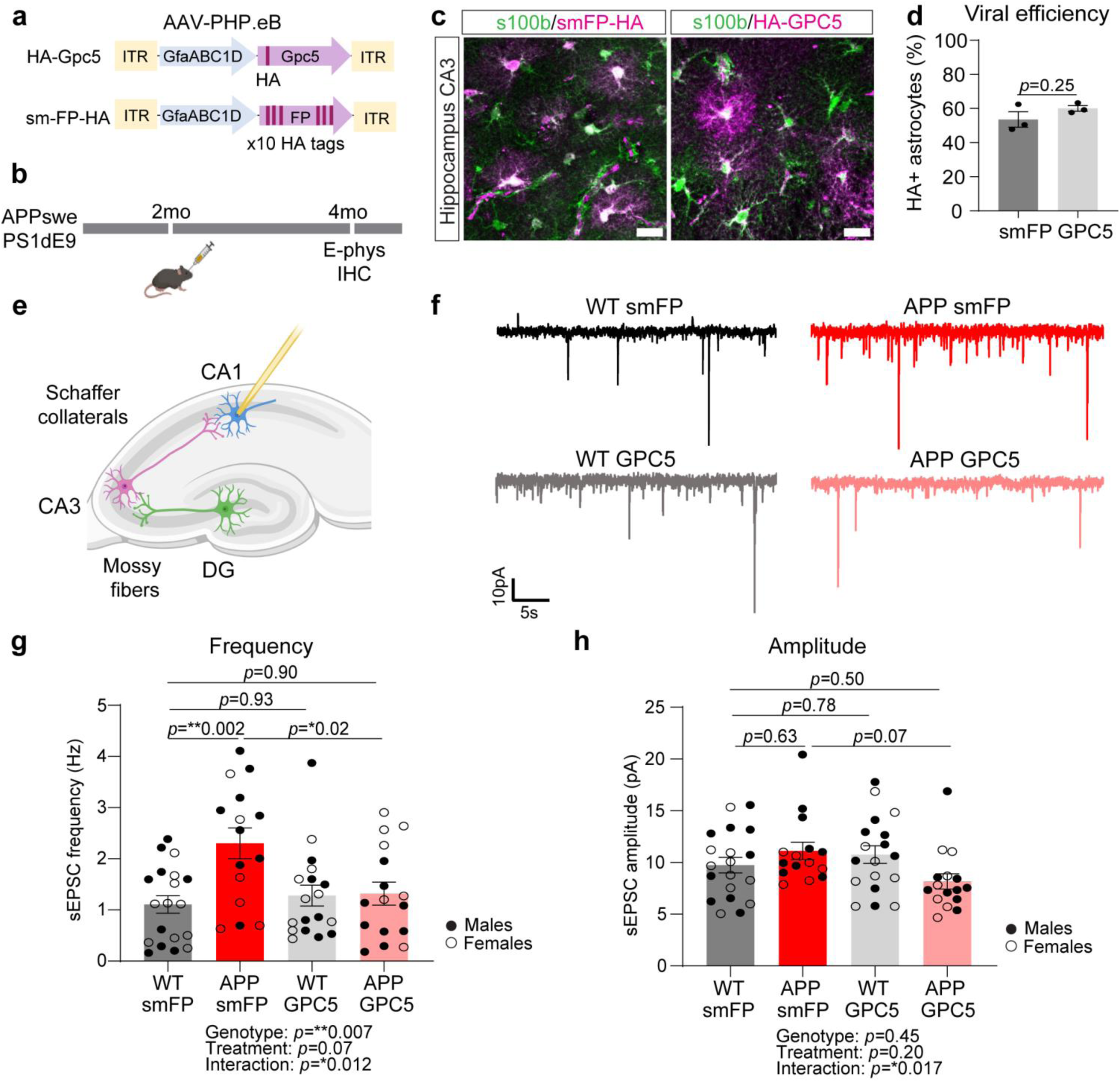
Astrocyte Glypican 5 overexpression prevents hippocampal synaptic hyperactivity in APP mice at 4 months. a-b) Diagram depicting a) AAV-PHP.eB expressing HA-Gpc5 or smFP as control under the astrocyte-specific minimal GFAP promoter; b) APP 2-month-old mice were retro-orbitally injected with AAV-HA-Gpc5 or AAV-smFP-HA as control, and at 4 months of age mice were collected for electrophysiology recordings. Same image is shown in Supplementary Fig 6. c-d) Representative image of 7-month-old hippocampi from mice injected with either HA-GPC5 or smFP-HA and stained with HA and s100b antibodies. Same images are shown in Supplementary Fig 6. Scale bar=20 µm. d) Quantification showing the percentage of s100b-positive astrocytes overexpressing the HA-Gpc5 or smFP-HA construct in the hippocampus. N=3 mice per group, statistical test: T test. e-h) Whole cell patch clamp recordings of spontaneous excitatory postsynaptic currents (sEPSC) in hippocampal CA1 pyramidal neurons. e) Diagram showing the hippocampal neurons recorded in CA1. f) Representative traces from different groups showing an increased frequency of sEPSC in the APP smFP group that is prevented APP GPC5 overexpressing group. g-h) Average frequency (g) and amplitude (h) of sEPSC events during 5-minute recordings. Each data point represents an independent neuron. n= WT smFP: 19 neurons, 9 mice, WT GPC5: 18 neurons, 10 mice, APP smFP: 15 neurons, 10 mice; APP GPC5: 16 neurons, 8 mice. Male (close circles) and female (open circles) mice were included for the analysis. Statistics: 2-way-ANOVA Tukey’s correction for multiple comparison run on neurons. * *p*<0.05, \*\**p*<0.01, \*\*\**p*<0.001. Graphs show the mean ± SEM.

To assess synaptic function, we performed *ex vivo* whole-cell patch clamp recordings of spontaneous excitatory postsynaptic currents (sEPSC) in hippocampal CA1 pyramidal neurons at 4 months (**Fig 6e**). This showed that APP-smFP mice have a significant increase in the frequency of sEPSCs relative to WT-smFP control, and that this increase does not occur in the APP-GPC5 group (**Fig 6f,g**). We also observed a significant increase in resting membrane potential in APP-GPC5 relative to APP-smFP (**Supplementary Fig 5d**), and an increased sEPSC decay time in both APP groups relative to WT, independent of GPC5 overexpression (**Supplementary Fig 5e**). Rise time and sEPSC amplitude are not changed across experimental groups (**Fig 6h, Supplementary Fig 5f**). In conclusion, we show here that overexpressing GPC5 in astrocytes *in vivo* is able to prevent early synaptic hippocampal hyperactivity in APP mice.

### 7. Astrocyte Glypican 5 overexpression improves spatial learning in APP mice at 6 months

To explore the functional consequences of GPC5 overexpression at the level of behavior, we assessed spatial learning and memory in APP mice at 6 months, that were injected with AAV.PHP.eB expressing HA-GPC5 or smFP control at 2 months (**Fig 7a**). Immunohistochemistry analysis confirmed that after 5 months of overexpression (7 months old, after behavioral testing) ∼50% of astrocytes in the hippocampus are targeted (**Fig 6c,d, Supplementary Fig 6**). We first tested locomotor capabilities using the open field test and found a significant increase in the distance travelled by APP mice relative to WT, which is not affected by GPC5 overexpression (**Supplementary Fig 7a**). This hyperactive phenotype has been previously reported in the same APP mouse model^45^. The speed of movement and time spent in the center of the arena are not different across groups (**Supplementary Fig 7b,c**). We then assessed spatial learning using the Barnes maze, with mice trained for 5 consecutive days (2 trials/day) to find an escape box in a circular arena using spatial cues (**Fig 7b**). We found that APP-smFP mice showed mild deficits in the rate of learning during the initial days of testing: while WT-smFP mice showed a significant reduction in the escape latency already after one day of training, APP-smFP did not. This deficit was prevented in the APP group overexpressing GPC5 (**Fig 7c-e**). We found no significant difference in the area under the escape latency curve across groups, nor in the performance during the probe trials, showing all groups learn the task after 5 days of training (**Supplementary Fig 7d,e**). During reversal learning, when the escape box is moved to the opposite hole, we found that GPC5 overexpressing mice displayed a significant reduction in the area under the escape latency curve regardless of genotype (2-way ANOVA treatment effect *p*=0.03) (**Supplementary Fig 7f,h**), with no changes in the learning rate (**Supplementary Fig 7i**). In summary astrocyte GPC5 overexpression prevented mild spatial learning deficits in APP mice and accelerated reversal learning in both WT and APP mice.

**Figure 7:**
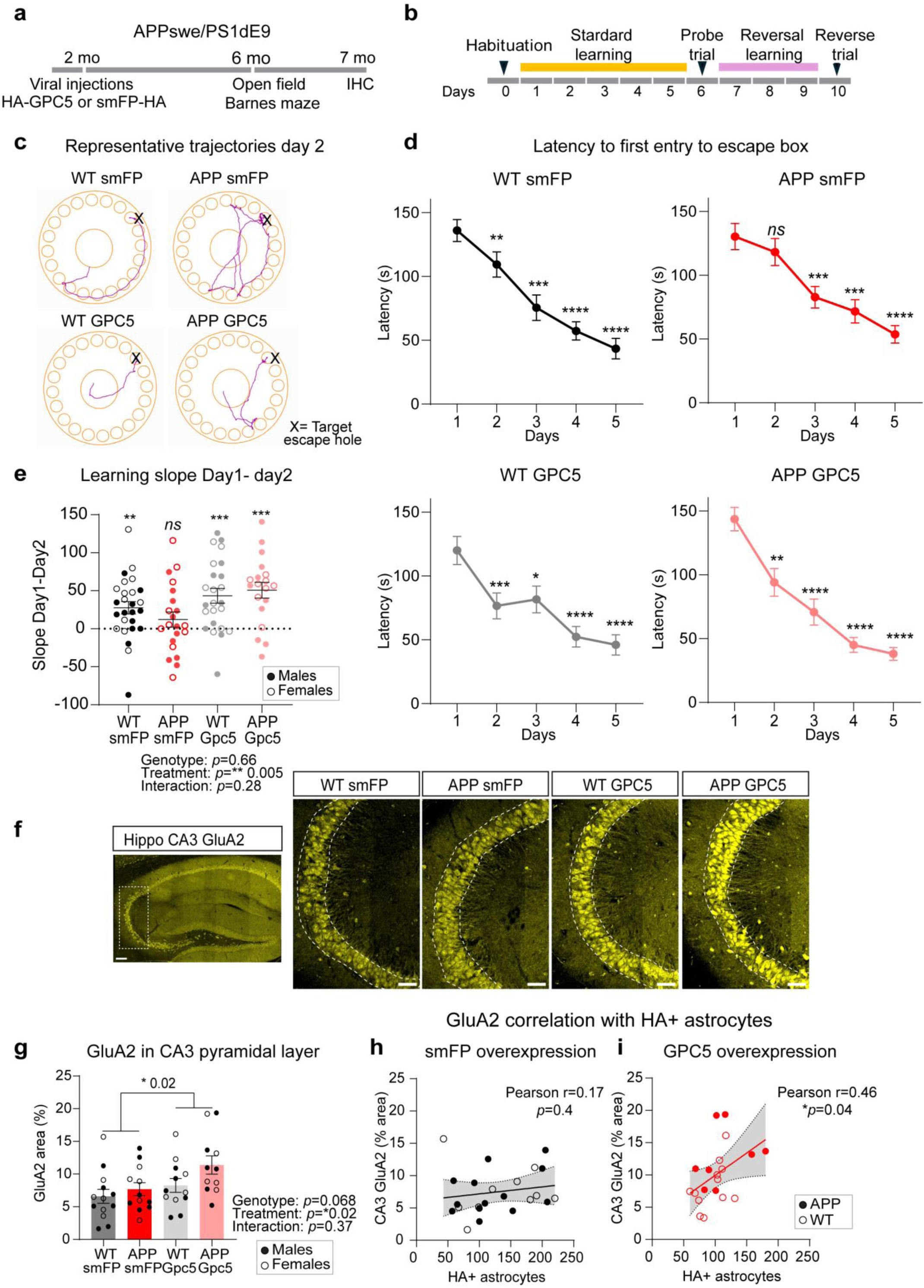
Astrocyte Glypican 5 overexpression improves spatial learning in APP mice at 6 months. a) APP and WT 2-month-old mice were retro-orbitally injected with AAV-HA-GPC5 or AAV-smFP as control. At 6 months of age mice were behaviorally tested in the open field and Barnes maze test, at 7 months brains were collected for immunohistochemistry analysis. b) Barnes maze memory test: mice were trained for 5 consecutive days (2 trials a day) to find an escape box. The next 3 days the escape box location is changed to the opposite hole to test cognitive flexibility. On days 6 and 10 a probe trial is performed when the box is removed. c) Representative trajectories used by mice to find the escape box during standard learning (day 2). d) Mean latency to find the target hole is plotted across different training days (mean of the 2 trials per day). Each panel shows different experimental groups. Statistics are calculated using 2-way ANOVA with Dunnett’s multiple comparisons test relative to day 1. WT smFP: N=27(13F, 14M); APP smFP: N=21 (10F, 11M); WT GPC5: N=24 (13F, 11M), APP GPC5: N=18 (7F, 11M). e) Learning slope between day 1 and day 2 (mean latency day 1-mean latency day 2). Statistical analysis is calculated using one-sample T test against hypothetical value of 0 and corrected for multiple comparisons. Each data point represents an independent mouse. (N same as in panel d). f-g) Representative tile scans stitched images of GluA2 immunostaining in the hippocampal CA3 region in 7-month-old APP and WT littermates overexpressing AAV-Gpc5 or smFP. Scale bar = 20 µm (f). Quantification of GluA2 coverage showed a significant increase in GPC5-overexpressing mice (2-way ANOVA treatment effect: * *p*= 0.02). WT smFP: N=13 (6F, 7M); WT GPC5: N=12 (6F, 6M); APP smFP: N=12 (5F, 7M); APP GPC5: N=11(5F, 6M). Scale bar= 100 µm. h-i) Pearson correlation between GluA2 area and number of smFP (h) or GPC5 (i)-overexpressing astrocytes showed a significant correlation (*p*=0.04) only in the GPC5 overexpressing group (i). Statistics: * *p*<0.05, ** *p*<0.01, *** *p*<0.001, **** *p*<0.0001

We then asked whether GPC5 overexpression impacted other AD-related pathologies in APP 7-month-old mice, collected after behavioral testing (**Fig 7a**). We found no significant differences in hippocampal amyloid plaque burden between APP-smFP and APP-GPC5 (**Supplementary Fig 8a-c**), or in GFAP immunoreactivity across all experimental groups (**Supplementary Fig 8d-f**). Astrocyte GPC5 impacts synapses by regulating presynaptic terminal size and postsynaptic recruitment of GluA2-containing AMPARs in the developing brain^22^. To ask if GPC5 mediates its effect on behavior through an impact on synapses, we characterized synaptic proteins using immunohistochemistry. We found a significant reduction in presynaptic vGlut1, but not synaptoporin, in CA3 *stratum lucidum* in 7-month APP mice relative to WT, with no impact of GPC5 overexpression (**Supplementary Fig 8g-l**). In contrast, we observed a significant upregulation of postsynaptic GluA2 AMPARs in GPC5-overexpressing groups compared to smFP control, regardless of genotype (2-way ANOVA treatment affect *p*=0.02, **Fig 7f,g**). We also found a significant positive correlation between the number of GPC5-overexpressing astrocytes and level of GluA2 (Pearson correlation 0.46, *p*=0.04), and this correlation is not present in smFP-overexpressing group (Pearson correlation 0.17, *p*=0.4) (**Fig 7h,i**). In summary, we show that *in vivo* astrocyte GPC5 overexpression is able to increase levels of GluA2 AMPARs in hippocampal CA3.

## DISCUSSION

In this study we have generated a comprehensive dataset characterizing sex and age-dependent astrocyte gene expression alterations in amyloid and tau pathology mouse models (**Fig 1**). We found that astrocytes show robust transcriptional alterations in the advanced stages of pathology, with mild sex differences, primarily in the APP model (**Fig 2**). Astrocytes in APP and Tau models show a progressive upregulation of immune-related genes, and a downregulation of genes important for synapse function such as *Gpc5* (**Fig 3, 4**). This signature is partially shared with human AD patients, suggesting that in AD brains astrocytes create an environment that allows synapse destabilization and dysfunction to occur (**Fig 5**). We show that manipulation of components of this signature by *in vivo* overexpression of GPC5 in astrocytes is sufficient to prevent early synaptic dysfunction and improve spatial learning in the APP mouse model (**Fig 6-7**).

An important novelty of our study is the inclusion of male and female mice for the assessment of sex differences. We found that very few genes showed a significant sex-genotype interaction, suggesting that sex does not strongly impact astrocyte DEGs in amyloidosis and tauopathy models at the ages we tested. This data is consistent with previous human AD postmortem studies which showed no major sex differences in the distribution of astrocyte disease-associated clusters in female relative to male patients^37^. However, we observed mild differences in the immune response in APP male mice, which are worth investigating in future experiments to understand how this may impact pathology progression.

Another important novelty of this report is the finding that tau pathology elicited greater changes in expression of GWAS AD risk genes in astrocytes compared to amyloid pathology (**Fig 4, Supplementary Fig 3**). This data contrasts with previous work showing that only amyloid pathology induced changes in AD risk genes in both astrocytes and microglia^14,46^. One explanation for this discrepancy is that the previous papers analyzed either less aggressive tauopathy mouse models^46^, or at earlier pathology stages (5 months) than our current study^14^, suggesting that long term exposure to tau pathology is needed to alter expression of GWAS AD risk genes in astrocytes. Notably, we found that astrocytes in both APP and Tau models upregulate AD risk genes normally enriched in microglia, including *Bcl3*, *Plcg2* and *Trem2.* TREM2 upregulation in astrocytes has also been shown in acute ischemic stroke^47^, and spinal cord injury^48^ models, suggesting it might be a generalized astrocyte response to pathology.

A goal of this study is to determine whether astrocyte synapse-related functions are altered in AD brains, and whether this contributes to AD synaptic dysfunction. We found that astrocytes in 12 month old APP and Tau mice show a downregulation of genes important for synapse regulation, which is shared with human AD, FTD and MS patients^35–38^ (**Fig 4**). Interestingly, this astrocyte signature is not exclusive to neurodegenerative disorders, and is also present in response to acute inflammatory stimulus^49^, and in a spinal cord injury mouse model^50^. This data suggests that astrocytes compromise their synapse supporting functions in response to different pathological triggers. Therefore, manipulating this signature in astrocytes might be beneficial not only in AD, but in a variety of different disorders.

Among the astrocyte genes that are significantly downregulated in APP and Tau models, and in human AD, MS, and FTD patients, is *GPC5*^35–38^. GPC5 belongs to the family of GPI-anchored heparan sulphate proteoglycans called glypicans, with 6 members in mammals (GPC1-6)^51^. GPC5 modulates synapse maturation and stabilization by regulating presynaptic terminal size and postsynaptic recruitment of GluA2-containing AMPARs^22^. We therefore hypothesized that GPC5 downregulation in neurodegenerative disorders may contribute to synaptic dysfunction. Using *ex vivo* patch-clamp electrophysiology we showed that *in vivo* astrocyte GPC5 overexpression is able to prevent the increased sEPSC frequency observed in CA1 neurons in 4-month APP mice (**Fig 6**). Additionally, GPC5 overexpression increases levels of GluA2 in the hippocampal CA3 in 7-month APP mice (**Fig 7**). A prediction is that GPC5-dependent recruitment of GluA2-containing AMPARs facilitates the strengthening of hippocampal CA3 synapses, and consequently enhances hippocampus-dependent spatial leaning in the Barnes maze test, as we showed (**Fig 7**).

Importantly, *GPC5* expression is not restricted to astrocytes and it is also expressed by oligodendrocyte progenitor cells, and some neurons. In humans, there are specific subtypes of excitatory and inhibitory neurons in the entorhinal cortex that express GPC5. Interestingly, GPC5-expressing excitatory neurons are specifically vulnerable to AD, and loss of these neurons strongly correlates with worse cognitive abilities^52^, suggesting that neuronal GPC5 might also be protective for synaptic function. Neuronal presynaptic GPC5 has been shown to bind to LRRTM4 receptor on the postsynaptic site, which in turn mediates synapse formation and AMPAR clustering^53^. In addition, other members of the glypican family have been shown to bind to presynaptic PTPRD and PTPRS^54,55^, and postsynaptic GPR158 receptors^56^. Future experiments should investigate if astrocytic GPC5 mediates its synaptic action by interacting with LRRTM4 or other candidate receptors in neurons.

Interestingly, *GPC5* downregulation occurs preferentially in astrocytes around amyloid and MS plaques and spinal cord injury lesions, environments that are highly pro-inflammatory^44,50^ (**Fig 5**). This data suggests that *GPC5* downregulation might be triggered by inflammatory signals. STAT3 is a transcription factor that can regulate the expression of GFAP and other reactive immune-response genes, and it has also been predicted to act as a GPC5 transcriptional regulator^48,57^. Future experiments should investigate if GPC5 downregulation is mediated by STAT3 or other transcription factors.

In conclusion, we provide here an in-depth profiling of astrocyte transcriptional alterations across different pathology stages in two different AD mouse models and in a sex-specific manner. This resource will allow for future experiments to understand the functional role of astrocytes in AD pathology progression. Using this dataset, we identified a novel astrocyte-specific mechanism contributing to synaptic dysfunction in an AD mouse model. We showed that *in vivo* manipulation of astrocyte GPC5, which we found downregulated in APP and Tau mouse models, is able to prevent early synaptic deficits in an amyloidosis mouse model, strengthen synapses and improve spatial learning. GPC5 downregulation is shared across different neurological disorders including FTD, MS and AD^13,35–38,43,44^. We propose that GPC5 downregulation is a common mechanism contributing to synaptic dysfunction in different neurodegenerative disorders, and that targeting GPC5 provides a new avenue for preventing synaptic alterations in multiple disorders.

## Acknowledgements

This work was supported by awards from the Chan Zuckerberg Initiative Neurodegeneration Network (DAF2018-191894), Coins for Alzheimer’s Research Trust (CART), Auen Foundation, Grohne Family Foundation and Hearst Foundation to N.J.A. I.H.S. was supported by an Alzheimer’s Association Research Fellowship (AARF-20-681410), BrightFocus Foundation Postdoctoral Fellowship Award (A20201645F), and the Anderson Foundation. This work was supported by the Waitt Advanced Biophotonics Core Facility of the Salk Institute (with funding from NIH-NCI CCSG: P30 CA01495, NIH NlA San Diego Nathan Shock Center P30 AG068635, and the Waitt Foundation); the GT3 Core Facility of the Salk Institute (with funding from NIH-NCI CCSG: P30 CA01495, an NINDS R24 Core Grant and funding from NEI (R24NS092943); the *In vivo* Scientific Service facilities; the Razavi Newman Integrative Genomics and Bioinformatics Core at the Sak Institute (with funding from NIH-NCI CCSG P30 CA014195, NIH-NIA San Diego Nathan Shock Center P30 AG068635, the NIH-NIA Liver Cancer P01 AG073084-04, the Howard and Maryam Newman Family Foundation and the Helmsley Trust); the Institute for Genomic Medicine (IGM) Genomics Center at UC San Diego supported by NIH SIG grant S10 OD026929; and the UC San Diego Shiley-Marcos Alzheimer’s Disease Research Center (with funding from NIA grant P30 AG062429). We thank the members of the Allen laboratory for providing feedback on the manuscript.

## Authors contribution

N.J.A. and I.S.H conceived of the project and designed the experiments. I.S.H. performed the experiments, designed the analysis pipelines, and conducted the data analysis. A.P. performed the electrophysiology recordings and data analysis. T.T performed the Western blot experiments and analysis. A.D. performed the immunostaining analysis for Tau line characterization. J.B. contributed to RNA sequencing analysis. C.S., M.M., S.M, F.A., T.D., A.F., contributed to immunostaining, image acquisition and analysis. Q.A. assisted with the behavioral studies. I.S.H. and N.J.A. interpreted the results and wrote the paper with comments from all authors.

## Materials & Correspondence

Correspondence and request for materials should be addressed to Nicola J. Allen or Isabel H. Salas

## Data and Code Availability

RNA sequencing sample information, gene expression level, DEG list and pathway analysis are available in supplementary tables. GEO accession will be available with the published manuscript.

## MATERIALS AND METHODS

### Mice

All animal experiments were approved by the Salk Institute Institutional Animal Care and Use Committee (IACUC). Mice were housed with a 12-hour light/dark cycle in the Salk Institute animal facilities. Access to food and water was provided ad libitum. Mice of both sexes were used for all experiments between the ages of 2 to 12 months.

### APPswe/PSEN1dE9

mice were obtained from Jackson Laboratory (MMRRC Strain #034832-JAX) and were crossed with Wild-type C57Bl6/J mice (Jax 000664) to obtain hemizygous litters. APPswe/PSEN1dE9 and WT littermates were used to characterize AD pathology at 4, 6 and 12 months (Fig 1) and for GPC5 overexpression experiments (Fig 6,7). This line is referred to throughout the manuscript as APP.

### MAPT P301S

mice were obtained from Jackson Laboratory (Jax Stock: 008169) in a B6;C3F1 background and were crossed with WT B6;C3 mice to obtain hemizygous litters. Tau P301S and WT littermates were used to characterize AD pathology at 4, 6 and 12 months (Fig 1). This line is referred to throughout the manuscript as Tau.

### APPswe/PSEN1dE9 x Aldh1l1-EGFP/Rpl10a

Aldh1l1-EGFP/Rpl10a mice were obtained from Jackson Laboratory (JAX:030247) and crossed with APPswe/PSEN1dE9. Mice expressing EGFP/Rpl10a and hemizygous for APPswe/PSEN1dE9 or WT were used for ribosome pulldown and RNA sequencing (Fig 1-5).

### Tau P301S x Aldh1l1-EGFP/Rpl10a

Aldh1l1-EGFP/Rpl10a mice (JAX:030247) were crossed with MAPT P301S. Mice expressing EGFP/Rpl10a and hemizygous for Tau P301S or WT were used for ribosome pulldown and RNA sequencing (Fig 1-5).

### Ribotag Pulldown

#### Tissue collection

4, 6, and 12-month-old Aldh1l1-EGFP/Rpl10a mice crossed with APPswe/PS1dE9 or Tau P301S were anesthetized by intraperitoneal injection of 100 mg/kg ketamine, 20 mg/kg xylazine. Brains were extracted, washed in ice cold PBS and dissected in ice cold 1x HBSS, 2.5mM HEPES-KOH (pH7.3), 35mM Glucose, 4mM NaHCO3, 100mg/ml cycloheximide. Hippocampus was microdissected, flash frozen in liquid nitrogen and stored at -80 until mRNA isolation. Tissue collection was performed between 2pm and 5pm for all samples.

#### Tagged ribosome pulldown and RNA extraction

Protocol adapted from^58,59^. Briefly, 2 hippocampi per mouse (each replicate = 1 mouse) were homogenized in 300µl of ice-cold dissection buffer (1% NP-40, 10mM KCl, 50mM Tris pH 7.4, 12mM MgCl2, 0.1mg/ml cycloheximide, 1mg/ml heparin, 1mM DTT, 1:200 Rnase IN added fresh), using a Pestle Motor (VWR 47747-370). After that, volume was brough up to 1ml with dissection buffer, and passed 10x through a 25G needle. Tissue homogenate was then centrifuged 10G for 10 mins at 4°C. 50µl of the supernatant was kept as “input” and the rest was incubated on a rotator at 4°C for 3 hours with 17µg of anti-GFP antibodies to bind the GFP-tagged ribosomes (HTZ-GFP-19C8, and HTZ-GFP-19F7 obtained from Memorial Sloan-Kettering Monoclonal Antibody Facility). The antibody-ribosome complex was then incubated overnight on a rotator at 4°C with magnetic IgG beads (Thermo Scientific Pierce #88847). Next day samples were washed 4 times with 800µl of high salt buffer 1% NP-40, 300mM KCl, 50mM Tris pH 7.4, 12mM MgCl2, 0.1mg/ml cyclohexamide, 1mg/ml heparin, 0.5mM DTT, Rnase IN. RNA was extracted using the Qiagen RNeasy Plus Mini kit (Qiagen 73404) according to manufacturer instructions, eluted in 30µl RNase-free water, and kept at -80°C until cDNA library preparation. RNA from the input pre-antibody incubation was extracted in parallel using the same procedure to determine astrocyte enrichment.

### RNA-sequencing and analysis

cDNA library preparation and RNA sequencing were run in 4 independent batches. All samples belonging to the same age (4, 6, 12 months) or mouse model (APP, Tau) were run in the same batch with the exception of Tau-12-month-old group which was run in 2 different batches. Batch 1: Tau 12-month-old samples (cohort 1); Batch 2: APP 6-months and 4-months; Batch 3: Tau 6-months and 4-months; Batch 4: APP 12-month-old and Tau 12-month-old (cohort 2). RNA concentration and quality were measured with a Tape Station (Agilent). mRNA was extracted with oligo-dT beads, capturing poly(A) tails, and cDNA libraries were made with the Illumina TrueSeq library. Stranded mRNA Library was prepared by the Institute for Genomic Medicine (IGM) Genomic center in University of California San Diego (UCSD). Samples were sequenced on an Illumina NovaSeq4 with paired-end 100 reads at 25 million reads per sample. FASTQ files were downloaded and quality-tested using MultiQC tool (v1.13). Alignment to the mm39 genome was performed using Spliced Transcripts Alignment to a reference (STAR) aligner, version 2.7.10a ^60^. Mapping was carried out using default parameters (up to 8 mismatches per paired read). MultiQC (v1.13) was used to summarize the uniquely mapped reads and other quality control parameters (data summarized in Supplementary Table 1). Samples with <20M reads assigned and <75% alignment were excluded from the analysis. Per-gene read counts were summarized using featureCounts (version 2.0.1)^61^ with default settings (not allowing multi overlaps, with minimum and maximum fragment length of 50 and 600bp respectively).

#### Differential gene expression analysis

Differential expression analysis was performed using DESeq2 R package (v1.38.3) with default normalization. The following multiple comparisons were performed independently for each age group (4, 6 and 12 months) and mouse model (APP and Tau). Outputs from these comparisons are shown in Supplementary Table 3,4 (Tg versus WT comparison, analysis 1, 2), and in Supplementary Table 7 (sex-genotype interaction, analysis 3).

- *Analysis 1*: Differential expression analysis was run based on sex and genotype combined (∼sex_genotype) and independent results for males and females were extracted resulting in two output groups: male transgenic versus male WT, female transgenic versus female WT. For Tau-12-month group, cohort was included as a variable (∼ Cohort + sex_genotype);

- *Analysis 2*: Differential expression analysis was run based on genotype and controlled for sex (∼Sex + Genotype). This resulted in one output group: Transgenic versus WT. For Tau-12-month group, cohort was included as a variable (∼Cohort + Sex + Genotype).

-*Analysis 3*: Differential expression analysis for sex-genotype interactions controlled for sex and genotype (∼Sex + Genotype + Sex:Genotype).

##### Criteria for establishment of differentially expressed genes (DEGs)

(Supplementary Table 5) Padj <0.05, mean transcripts per million (TPM) across genotypes TPM >1, Log2FC > |0.3|

##### Gene-set enrichment/Over-representation analysis

(Fig 2, 4). For gene set enrichment analysis genes were ranked based on Log2 fold changes and the webgestalt website (https://www.webgestalt.org) or fgsea package in R (v1.24.0) was used to run the analysis. Affinity propagation tool from webgestalt website was used to select non-redundant significant reactome pathways enriched.

##### Time-course analysis of DEGs

(Fig 3) 12-month-old DEGs were selected and responses were categorized based on the LFC at 6 and 12-month-old, and significance status at 12 months, but independently of significance status at 6 months.

- Up-Up response: 6 months: LFC>0.3; 12-month-old LFC>0.3, *padj* <0.05

- Unchanged-Up response: 6 months: -0.3<Log2FC<0.3; 12-month-old: LFC>0.3, *padj* <0.05

- Down-Up response: 6 months: LFC<-0.3; 12-month-old LFC>0.3, *padj* <0.05

- Down-down response: 6 months: LFC<-0.3; 12-month-old LFC<-0.3, *padj* <0.05

- Unchanged-down response: 6 months: -0.3<Log2FC<0.3; 12-month-old: LFC<-0.3, *padj* <0.05

- Up-down response: 6 months: LFC>0.3; 12-month-old LFC<-0.3, *padj* <0.05

##### Astrocyte synapse-regulating gene list generation

(Fig 4) Astrocyte synapse-regulating gene sets were created with the following criteria:

- Synaptogenic and synapse-eliminating factors: Gene list was based on previous literature, with a mean TPM >10 using the WT 12-month-old dataset

- GABAergic synapse: All genes with a mean TPM >10 in the WT 12-month-old dataset, that contained the name GABA in the gene description and were involved in GABA transport, reception of synthesis.

- Glutamatergic synapse: All genes with a mean TPM >10 in the WT 12-month-old dataset, that contained the name glutamate in the gene description and were involved in glutamate transport, reception or synthesis.

- Potassium channels: All genes with a mean TPM >10 in the WT 12-month-old dataset, that contained the name potassium in the gene description.

##### AD risk genes astrocyte enrichment

(Fig 4) List of AD risk genes were obtained from 3 different GWAS reports^32–34^. To determine the AD risk genes that were expressed or enriched in astrocytes, the following criteria was used:

- Expressed in astrocytes: Mean TPM >1 using the WT 12-month-old dataset from the APP group.

- Enriched in astrocytes: WT from 12-month-old dataset was used to run differential expression analysis comparing pulldown versus input samples (∼sampleType). DEGs with *padj* <0.05, Log2FC pulldown/input > 1 were considered significantly enriched in astrocytes.

##### Human-mouse correlation analysis

(Fig 5) Ensembl Biomart (ensembl.org/info/data/biomart/index.html) was used to identify mouse orthologous genes from human astrocytes DEGs identified in Grubman 2019 dataset ^35^. The LFC of the human DEGs was plotted against the LFC from 12-month-old mice (APP and Tau). Overlapping DEGs were plotted using VennDiagram package in R (v1.7.3).

### Western blot analysis for synaptic markers

#### Tissue collection

4, 6 and 12-month-old APP and Tau mice were anesthetized by i.p injection of 100 mg/kg ketamine, 20 mg/kg xylazine. Hippocampi were rapidly dissected, snap frozen in liquid nitrogen and stored at -80⁰C until use. Hippocampi were lyzed in 300 µL of RIPA buffer (ThermoFisher, cat#89900) supplemented with proteinase inhibitors (Thermo Scientific 1861278) and phosphatase inhibitors (Thermo Scientific 1862495) and incubated for 30 minutes at 4⁰C with rotation. Samples were then centrifuged at 12000 rpm for 10 minutes at 4⁰C, the supernatants collected, and the protein concentration quantified using Bradford assay. Lysates were diluted to 2 µg/µL after supplementing with the 5x reducing sample buffer (ThermoFisher, Cat#39000), incubated at 55 ⁰C for 45 min, and stored in -20 ⁰C until running the Western blot. 10 µL (20 µg total protein) samples were loaded on the blot.

#### Western blot

10 µL (20 µg total protein) samples were separated on 4-12% Bis-Tris gradient gels (Invitrogen NW04122BOX) at 150V for 1 hour in MOPS SDS buffer (Invitrogen B0001.02). Proteins were transferred to PVDF membrane (Cytiva #10600022) at 80V for 2 hours in an ice bucket using Tris-Glycine buffer (Thermo Fisher Scientific 28363) with 20% methanol (Fisher A412-4). The membrane was cut to separate the sections with the molecular weight of interest, blocked in blocking buffer containing 1% Casein in TBS (Biorad #1610782) for 1 hour at room temperature and incubated overnight at 4⁰C in blocking buffer with primary antibodies. The next day the membrane was washed for 3×10 minutes with TBS with 0.1% Tween-20 and incubated with secondary antibodies diluted in blocking buffer for 1 hour at room temperature. The membrane was then washed 3×10 minutes with TBS with 0.1% Tween-20, bands were visualized using the Odyssey Clx Infrared Imager (Li-Cor) and quantification of band intensity was performed using ImageStudio software (Li-Cor). Based on their MW and antibody hosts, PSD-95 (mouse), Synapsin (rabbit), Synaptophysin (rabbit) were blotted from gel 1, GluA1 (rabbit) and vGlut1 (guinea pig) from gel 2, GluA2 (rabbit) and vGlut2 (rabbit) from gel 3, each with its own β-actin (mouse) loading control. DyLight 800 for PSD-95 and β-actin, and AlexaFluor-680 for others.

#### Antibodies

| Primary Antibody | Species | Catalog # | Dilution |
| --- | --- | --- | --- |
| GluA1 | Rabbit | Millipore #AB1504 | 1:1000 |
| GluA2 | Rabbit | Millipore #AB1768-I | 1:1000 |
| Synaptophysin | Rabbit | Abcam #ab16659 | 1:1000 |
| Synapsin | Rabbit | Millipore #574778 | 1:1000 |
| vGlut1 | Guinea pig | Millipore#AB5905 | 1:3000 |
| vGlut2 | Rabbit | Synaptic systems#135403 | 1:1000 |
| PSD-95 | Mouse | NeuroMab #75-028 | 1:1000 |
| β-actin | mouse | Sigma Aldrich#A1978 | 1:5000 |
| Secondary Antibody | Emission | Catalog # |  |
| Goat anti-Mouse | DyLight 800 4X PEG | ThermoFisher #SA535521 | 1:20000 |
| Goat anti-Rabbit | Alexa Fluor 680 | ThermoFisher#A21109 | 1:20000 |
| Goat anti-Guinea Pig | DyLight 680 | ThermoFisher#SA5-10098 | 1:20000 |

### Cloning and AAV-PHP.eB packaging

For the overexpression of HA-tagged GPC5, cDNA for the coding sequence of mouse GPC5 (Origene # MR218095) was used. HA tag was cloned 72 bp (24aa) downstream of the N-terminus to avoid interference with the endogenous signaling peptide. This was achieved by cloning two different fragments of HA-GPC5 sequence: the N-terminal fragment containing the endogenous signaling peptide upstream the HA tag and the InFusion sequence optimized for InFusion cloning (Takara Bio #638909) was synthetically generated by IDT (TACGACTCACTATAGGCTAGCGCCACCATGGATGCCCGCACCTGGCGTCTGGGCTGGCGCTGTCTCCTTCTCCTGGCTCTCCTTGGATCTACCCGAAGCTATCCGTATGATGTTCCGGATTATGCAGAGGGCGTGG). The C-terminus fragment containing the rest of the mouse GPC5 sequence was first amplified using the following primers (forward: GAGGGCGTGGAGAGCTGCGA, reverse: TTACCAAAGCCAGGGAAGAAACATCAGTGT), and then amplified again with primers for inFusion cloning (forward: GGATTATGCAGAGGGCGTGGAGAGCTGCGAA, reverse: CATGTCTGCTCGAAGCGGCCGCTTACCAAAGCCAGGGAAGAAACATC). Both linearized fragments were run out on a 0.8% agarose TAE gel and extracted. We used a pZac2.1 backbone for overexpression. pZac2.1-gfaABC1D tdTomato (GFAP-tdTomato) from Addgene (#44332) was digested with NheI and NotI to remove the tdTomato CDS. The linearized product was run out on a 0.8% agarose TAE gel and extracted. An InFusion reaction was performed with the two linearized HA-GPC5 fragments and the linearized vector. Clones were selected using carbenicillin and sequenced to confirm presence of the inserted HA-GPC5. The cloned HA-GPC5 plasmid was validated in vitro using astrocyte cell cultures. As control we used the smFP-HA vector (#59759) cloned under the same gfaABC1D minimal promoter as previously described in ^62^. Both HA-GPC5 and smFP-HA plasmids were packaged into an AAV-PHPe.B virus by the Salk Institute Gene Transfer, Targeting, and Therapeutics Core and was.

### Mouse immunohistochemistry analysis

#### Tissue collection and sectioning

Mice were anesthetized by intraperitoneal injection of 100 mg/kg ketamine /20 mg/kg xylazine (Anased) mix and transcardially perfused with PBS followed by 4% paraformaldehyde (PFA) (Electron Microscopy Sciences 50980487). Brains were dissected and postfixed overnight in 4% PFA at 4°C. Next day brains were washed in PBS and incubated in 30% sucrose for 3-5 days at 4°C. Brains were then embedded in TFM (Tissue Freezing Media, Electron Microscopy Sciences 72593) frozen in a dry ice/ethanol slurry solution and stored at -80°C until use. Brains were sectioned in 16–20 µm-thick sagittal sections using a cryostat (Hacker Industries #OTF5000) and mounted on Superfrost Plus slides (Fisher #1255015). At least 3 mice were used for each experimental group and 2-3 sections were imaged and analyzed.

#### Immunostaining

Brain sections were placed in a humidified chamber to be blocked and permeabilized in blocking buffer (5% goat serum in PBS with 0.3% Triton X-100) for 1 hour at room temperature. Sections were then incubated over night at 4°C with primary antibodies diluted in the same blocking buffer (see list of antibodies below). Next day sections were washed for 3×5 minutes with PBS and incubated for 2 hours at room temperature with secondary antibodies diluted in blocking buffer (antibodies listed below). Then sections were washed again for 3×5 minutes in PBS, incubated for 10 minutes with DAPI (Thermo Fisher Scientific 62247) (1:5000), mounted with Fluoromount G (Southern Biotech 0100-01) and covered with 1.5mm-thick coverslip. For 6e10 antibody immunostaining, antigen retrieval was performed prior to blocking by 20 minutes incubation with 10 mM sodium citrate (pH 6.0) at 90°C.

#### Antibodies for immunohistochemistry

| Primary Antibody | Species | Company and Catalog # | Dilution |
| --- | --- | --- | --- |
| Synaptoporin | Rabbit | Synaptic systems 102002 | 1:500 |
| GluA2 | Rabbit | EMD Millipore AB1768 | 1:500 |
| vGlut1 | Guinea pig | EMD Millipore ab5905 | 1:2000 |
| 6e10 | Mouse | Biolegend 803015 | 1:500 |
| OC | Rabbit | Rockland 200-401-E87 | 1:500 |
| AT8 | Mouse | ThermoFisher mn1020 | 1:500 |
| S100b | Rabbit | Abcam AB52642-1001 | 1:500 |
| GFAP | Rabbit | Abcam AB7260 | 1:500 |
| GFAP | Mouse | EMD Millipore mab 360 | 1:500 |
| HA | Rat | Roche 3F10 | 1:100 |
| NeuN Alexa 488 | Rabbit | Abcam AB190195 | 1:500 |
| NeuN Alexa 647 | Rabbit | Abcam AB190565 | 1:500 |
| GPC5 | Rabbit | ThermoFisher PA5-83490 | 1:100 |
| GFP | Chicken | Millipore 06-896 | 1:100 |
| **Secondary Antibody** | **Species** | **Catalog #** | **Dilution** |
| Alexa 594 | Goat-anti-Rabbit | ThermoFisher A11037 | 1:500 |
| Alexa 594 | Goat-anti-Mouse | ThermoFisher A21424 | 1:500 |
| Alexa 488 | Goat-anti-Guinea pig | ThermoFisher A11073 | 1:500 |
| Alexa 594 | Goat-anti-Rat | ThermoFisher A11007 | 1:500 |
| Alexa 488 | Goat-anti-Chicken | ThermoFisher A11039 | 1:500 |

- AD pathology characterization: (Supplementary Fig 1)

APP: 6e10 mouse, GFAP Rabbit, Goat-anti-Mouse Alexa 594, Goat-anti-Rabbit Alexa 488 Tau: Staining 1: AT8 Mouse, NeuN-488 Rabbit, goat-anti-Mouse Alexa 594 Statining 2: GFAP Rabbit, goat-anti-Rabbit Alexa 594

- BacTrap line characterization: (Supplementary Fig 2)

GFP chicken, s100b rabbit, goat-anti-chicken Alexa 488, roat-anti-Rabbit Alexa 594 GFP chicken, NeuN-647 rabbit, goat-anti-chicken Alexa 488

- Viral efficiency characterization: (Supplementary Fig 6)

S100b rabbit, HA rat, goat-anti-rabbit Alexa 488, goat-anti-rat Alexa 594 NeuN-488 rabbit, HA rat, goat-anti-rat Alexa 594 GPC5 rabbit, goat-anti-rabbit Alexa 594

-Synapse characterization: (Fig 7, supplementary Fig 8)

Vglut1 guinea pig, GluA2 rabbit, goat-anti-rabbit Alexa 594, goat-anti-guinea pig Alexa 488 Synaptoporin Rabbit, vGlut1 guinea pig, goat-anti-rabbit Alexa 594, goat-anti-guinea pig Alexa 488

- Plaques/astrogliosis characterization: (Supplementary Fig 8)

GFAP mouse, OC Rabbit, goat-anti-Rabbit Alexa 594, goat-anti-mouse Alexa 488

#### Image acquisition and analysis

Imaging was performed using an Apotome Axio Imager Z2 epifluorescent microscope (Zeiss) AxioCam HR3 camera at 20x magnification. Tile images that contained the entire hippocampus were acquired for all experiments except for synapse analysis (vGlut1, GluA2, synaptoporin), where tiles were restricted to the CA3 hippocampal region. Images were acquired as 14-bit images, 0.233×0.232×1µm pixel size, as a z-stack with a total thickness of 7-8µm depending on the experiment. Imaging settings (lamp intensity, exposure time) were maintained across groups for each experiment. All image analysis was performed using FIJI as single-plane maximum projection images.

##### GFAP quantification

Entire hippocampus (Supplementary Fig 8) or CA3 region (Supplementary Fig 1) were selected as a region of interest (ROI) using the GFAP channel and images were thresholded equally across all experimental groups. Thresholded GFAP area normalized to ROI area was plotted as GFAP% area.

##### Amyloid plaque quantification

(Supplementary Fig 8) Entire hippocampus was selected as ROI and imaged was thresholded equally across all experimental groups using the OC channel. FIJI plug-in “Analyze Particles” was used to identify plaques with a minimum area of 50 µm^2^. Plaque area was plotted normalized to hippocampal ROI area.

##### GluA2 quantification

(Fig 7) Same size CA3 ROI was selected and GluA2 channel was thresholded equally for all experimental groups. To analyze the GluA2 signal in the CA3 cell body layer we used the Analyze Particles plug-in to identify cell bodies with a minimum area of 20 µm^2^. Thresholded GluA2 area within these particles normalized to the ROI selected was plotted as % area.

##### vGlut1 quantification

(Supplementary Fig 8) CA3 *stratum lucidum* was manually delimited as an ROI. Thresholded vGlut1 area within the *stratum lucidum* was quantified normalized to total *stratum lucidum* ROI area and plotted as % area. S*tratum lucidum* ROI area was not significantly changed across experimental groups (data not shown).

##### Synaptoporin quantification

(Supplementary Fig 8) CA3 ROI was selected and synaptoporin channel was thresholded equally for all experimental groups. Thresholded synaptoporin area normalized to the ROI area and plotted as % area.

##### Aldh1l1-EGFP/Rpl10a line characterization

(Supplementary Fig 2) Percentage of GFP+ /s100b+ astrocytes and GFP+ /NeuN+ neurons were quantified manually in the CA1 hippocampal region from 6-month-old Aldh1l1-EGFP/Rpl10a mice.

##### Viral overexpression characterization

(Supplementary Fig 5) Percentage of HA+/s100b+ astrocytes and HA+/NeuN+ neurons was quantified manually in the CA3 hippocampal region for WT mice overexpressing either GPC5 or smFP. Mice with no astrocytes infected were excluded from the analysis.

### Single-molecule RNA fluorescence *in situ* hybridization (smFISH)

smFISH was used to validate the dysregulation of *Gpc5*, *Bcl3* and *Hbegf* expression in 12-month-old APPswe/Psen1dE9 astrocytes. FISH was performed with the RNAScope Fluorescent Multiplex V2 Detection kit (ACD 323110) according to the manufacturer’s instructions for fixed experiments, followed by immunostaining of the same tissue. APPswe/Psen1dE9 12-month-old brains were collected and sectioned as described for immunohistochemistry analysis. Each probe was hybridized on different brain sections (2-3 sections per mouse, at least 6 mice per experiment). For smFISH slides were first baked for 30 mins at 60°C, postfixed in 4%PFA and dehydrated with increasing ethanol concentrations (50-70-100%). Sections were treated for 10 minutes with hydrogen peroxide and antigen retrieval performed by incubation with 10 mM Sodium citrate at 90°C for 25 minutes. After 3×5 minutes washes with PBS slices with incubated with protease IV in the HybEZ oven for 30 mins at 40°C, followed by 2 hours hybridization at 40°C with the appropriate probes: *Bcl3* (ACDbio 528431), *Hbegf* (ACDbio 437601), *Gpc5* (ACDbio 442831). Amplification steps were performed as instructed by manufacturer and signal was developed using HRP-C1 followed by incubation with Opal 570 (Akoya FP1488001KT) with the following dilutions: *Bcl3* 1:750, *Hbegf* 1:1000, *Gpc5* 1:1500. Reaction was blocked by incubation with HRP blocker and immunostaining was immediately started by 1 hour blocking in 10% normal goat serum, 0.2% triton X-100. Primary antibody incubation was performed over night at 4°C in a humidified chamber and the antibodies were diluted in antibody buffer (5% normal goat serum, PBS, 0.2% Triton X-100) (primary antibodies: Rabbit s100b, abcam, AB52642-1001, 1:500, Mouse 6e10 Biolegend 803015, 1:500). Next day sections were washed 3×5 minutes with PBS and incubated with secondary antibodies for 2 hours at room temperature diluted in antibody buffer (Goat-anti-Rabbit Alexa 488 ThermoFisher A11034, 1:500, Goat-anti-Mouse Alexa 647, ThermoFisher 21058). Then sections were washed again for 3×5 minutes in PBS, incubated for 10 minutes with DAPI diluted 1:5000 in PBS, mounted with Fluoromount G (Southern Biotech 0100-01) and covered with 1.5-mm-thick coverslip.

#### Image acquisition and analysis

##### *Bcl3* and *Hbegf* analysis

(Supplementary Fig 3) Images were acquired on a Zeiss LSM-880 confocal microscope using a 40x oil immersion objective. 4-tile images in the DG-CA3 hippocampal region were obtained as 8-bit, 0.21µm x 0.21µm x 1.34µm pixel size and as a z-stack of 7 slices with a total thickness of 8µm. Imaging settings were maintained across groups for each experiment. Cell profiler software (4.2.6) was used to quantify the area of smFISH probes expressed in s100b-positive astrocytes using maximum projection images. First, astrocytes and *Bcl3* or *Hbegf* puncta were identified as primary objects by using otsu and global thresholding methods respectively. Then, using the relate objects module, each probe puncta was assigned to their parent astrocyte. Data was extracted in excel and Matlab was used to calculate the total probe area expressed per astrocyte. The average *Bcl3* or *Hbegf* area in WT or APP astrocytes expressing the probe was plotted using Prism (version 8.4.3)

##### *Gpc5* analysis

(Fig 5) Imaging was performed using an Apotome Axio Imager.Z2 fluorescent microscope (Zeiss) AxioCam HR3 camera (Zeiss) at 20x magnification focusing on the DG-CA3 hippocampal region. Images were acquired as 14-bit images, 0.233 x 0.232 x 1µm pixel size, as a z-stack of 8 slices with a total thickness of 7µm. Imaging settings were maintained across groups for each experiment. Cell profiler software (4.2.6) was used to first identify plaque-associated and non-plaque associated astrocytes, and then calculate the area of Gpc5 probe expressed in each astrocyte. For this, amyloid plaques and s100b-positive astrocytes were first identified as primary objects using the global or sauvola thresholding methods respectively. Then, astrocytes within 7µm distance to amyloid plaques were identified using the relate objects module and classified as plaque-associated astrocytes. Then Gpc5 puncta was identified using global thresholding methods, and assigned to each astrocyte subtype using the relate objects module. Data was extracted in excel and Matlab and used to calculate the total Gpc5 area expressed per astrocyte. Then Gpc5 area per astrocyte was normalized to astrocyte area and was plotted as % area using Prism.

#### Human postmortem brain section immunostaining

Paraffin-embedded thin sections of postmortem human brain tissue from deidentified donors were obtained from the UC San Diego Shiley-Marcos Alzheimer’s Disease Research Center neuropathology core (funded with NIA grant P30 AG062429). All participants consented to brain donation at the time of enrollment in the ADRC. Sections were deparaffinated by baking at 60°C for 35 mins, followed by a xylene treatment (2×5 minutes), and rehydration using decreasing ethanol concentration washes (100-95-50-20%). Sections were then washed in 0.1%Tween-PBS, permeabilized for 10 minutes in PBS 0.3% triton, and incubated with 1× TrueBlack in 70% ethanol for 5 minutes. Antigen retrieval was performed by incubation with 10 mM sodium citrate (pH 6.0) at 90°C for 10 minutes and sections were blocked for 1 hour in blocking buffer (5% normal goat serum, 2% BSA in PBS), and incubated overnight with primary antibodies diluted in same blocking buffer. Primary antibodies: Rabbit anti-GPC5 (Invitrogen PA5-83490, 1:100), Rat anti-GFAP (ThermoFisher 130300, 1: 300). Next day sections were washed for 3×10 minutes in PBS, and incubated for 2 hours with secondary antibodies diluted in blocking buffer (secondary antibodies: AlexaFluor 555-goat anti Rabbit IgG, (ThermoFisher A-21428, 1:500), AlexaFluor 647 goat anti Rat IgG(ThermoFisher A-21247, 1:500). Sections were washed for 3×10minutes, DAPI treatment and mounting was performed as described previously. Imaging was performed in layer 1 of frontal cortex with a 20x objective on a Zeiss LSM700 confocal microscope. Images were acquired as 8-bit images, 0.31µm x 0.31µm x1µm pixel size, as a z-stack with a total thickness of 7µm depending on the experiment. Example image shown corresponds to a Braak stage 6 female AD patient.

#### Electrophysiology

##### Acute slice preparation

Coronal brain slices were prepared from 4-month-old animals injected with HA-GPC5 or HA-smFP. Animals were deeply anesthetized by i. p. injection with a 2.5% Avertin solution (0.4 g/kg). The brain was removed and cut into 300 µm coronal sections using a Leica VT1000s vibratome. The brain dissection was performed at physiological temperature (37⁰C), with choline chloride-based dissection solution composed of (in mM): 93 choline chloride, 2.5 KCl, 2 MgCl_2_, 2 CaCl_2_, 1.2 NaH_2_PO_4_, 20 HEPES, 5 Sodium L-ascorbate, 25 D-glucose and 24 NaHCO3. Slices were then placed in a recovery chamber containing HEPES artificial cerebrospinal fluid (aCSF) composed of (in mM): 5 HEPES, 93 NaCl, 2.5 KCl, 2 MgCl_2_, 2 CaCl_2_, 1.2 NaH_2_PO_4_, 5 Sodium L-ascorbate, 25 D-glucose and 24 NaHCO3. Slices recovered for 10 min at 37°C and then for at least 50 min at room temperature before recordings were performed for 4-6 hours after slicing. Solutions were supplemented with 1 mM sodium pyruvate and continuously equilibrated with carbogen (95% O2/ 5% CO2).

##### Electrophysiology recordings

Slices were placed in a recording chamber and perfused with carbogen-saturated aCSF composed of (in mM): 126 NaCl, 2.5 KCl, 1.2 MgCl_2_, 2.5 CaCl_2_, 1.2 NaH_2_PO_4_, 11 D-glucose and 25 NaHCO3, supplemented with 1 mM sodium pyruvate. Whole-cell recordings were performed from the soma of visually identified CA1 pyramidal neurons using infrared camera on a Scientifica microscope. Patch pipettes (thin-walled borosilicate glass pipette; Harvard Apparatus #BS4 64-0805; 4-6 MΩ of resistance) were filled with an internal solution composed of (in mM): 105 K-gluconate, 30 KCl, 10 phosphocreatine, 10 HEPES, 4 ATP-Mg, 0.3 GTP-Tris, 0.3 EGTA and 0.2% biocytin, adjusted to pH 7.2 with KOH. Recordings were performed using Axon Digidata 1550A and a Multiclamp 700B amplifier (Molecular Devices). All recordings were sampled at 10 kHz. Spontaneous excitatory postsynaptic currents (sEPSCs) were recorded at a membrane holding potential of –60 mV. All currents were recorded at room temperature (22–24⁰C), and only one neuron was studied in each slice. After a stabilization period of 10 min, sEPSCs were recorded for 5 minutes. Recordings were discarded if the access resistances were > 20 MΩ or changed > 25% during the recording. Electrophysiology recording analyses were performed using MiniAnalysis (Synaptosoft) and sEPSCs were detected using a 1 kHz post-hoc filter.

##### Acute slice immunostaining and imaging

After recording slices were transferred to a freshly prepared solution of PBS with 4% paraformaldehyde (PFA) and fixed overnight at 4⁰C. Next day sections were washed 3×10 minutes with PBS, blocked with 10% NGS, 0.3% triton in PBS and incubated over night at 4C with primary antibodies diluted in antibody buffer (3% NGS, 0.3% triton in PBS). Primary antibodies: Rabbit anti-HA 1:500 (Cell signaling 3724S), Mouse anti-GFAP 1:500 (mab 360). On day 2 sections were washed 3×10 minutes with PBS and incubated for 2 hours at room temperature with secondary antibodies diluted in antibody buffer. Secondary antibodies: Alexa anti-Rabbit-488 1:500 (Invitrogen A11034), Alexa anti-mouse 674 1:500 (Invitrogen 21058). After that sections were washed 3×10 minutes, incubated for 10 minutes with DAPI diluted 1:5000 in PBS, mounted with Fluoromount G (Southern Biotech 0100-01) and covered with 1.5mm thick coverslip.

##### Image acquisition and analysis

GFAP and HA co-stained images were acquired on a Zeiss LSM-880 confocal microscope with the 20x objective in CA1 and CA3 hippocampal regions as 8-bits, 0.42µm x 0.42µm x 1.12µm pixel size and as a z-stack of 21 slices with a total thickness of 22.492µm. Imaging settings were maintained across groups for each experiment. GFAP quantification was performed using FIJI as described in the previous immunohistochemistry section.

#### Mouse Behavior

Behavioral testing was performed in the Salk Institute’s Behavioral Testing Core (BTC) in the following order: Open field, Barnes maze. All tests were performed in the afternoon between the hours of 1pm and 5pm. Behavioral assays were run in 4 independent cohorts that were balanced for sex, genotype and factor being overexpressed. All behavioral tests were performed blinded to genotype and condition. Male mice were always tested before females. All mice were habituated to the test room for at least 30 minutes before the beginning of the test. Brain tissue was collected after the end of the behavioral testing.

##### Open Field

The test consists of a rectangular arena 40.6 x 40.6 x 38.1 cm; Med Associates Inc. ENV-515S-A with transparent plexiglass wall. The room was lit by indirect lighting of ∼30 lux. During the test, mice were placed in the center of the arena and allowed free exploration for 10 minutes. After that, mice were returned to their home cage and placed back into the housing facility. Activity Monitor software (Med Associates) was used to assess the locomotor activity. The total distance traveled, average speed and time in the center was assessed.

##### Barnes maze

The Barnes maze is a circular white table with 92 cm diameter elevated 1 m above the ground. Around the periphery, there are 20 holes with a diameter of 5 cm at a regular interval. A rectangular plastic box of 11.5cm x 6.5cm x 6cm deep was attached underneath one hole of the maze and used as the escape chamber. Four distal cues were placed ∼30 cm from the edge of the maze at regular intervals. Cues were roughly 50 cm2 and were made of cardboard and colored paper, resulting in different shapes and colors. Mice were habituated to the testing room for 30 minutes prior to initiation of the test. To overcome orientation bias, at the start of each trial the mouse was placed in the center of the maze and then covered with an opaque cylinder for 15 seconds before being allowed to explore the maze. The location of the escape chamber was kept constant for each mouse, but altered between mice. AnyMaze software was used to record and track the trajectory of the mice throughout all trials and habituation. The complete Barnes maze test consisted of 11 consecutive days; day 0 habituation (1 minute), day 1–5 acquisition trials (2 trials/3 minutes per day), day 6 probe trial (90 seconds), day 7–9 reversal trials (2 trials/3 minutes per day) and day 10 reverse probe trial (90 seconds). <u>Habituation</u>: mice were allowed to freely explore the maze for 1 minute. If the mouse did not enter the escape chamber within 1 minute, it was gently guided towards the escape hole, and the mouse was left there for 30 seconds, before returning to its home cage. <u>Acquisition trials</u>: mice were allowed to explore the maze for 3 minutes. When mouse entered the escape box, video was stopped, and mouse was returned to its home cage. If by the end of the test the mouse had not entered the escape chamber, it was gently guided towards the target hole and escape latency was set at 180 seconds. After all male mice were tested ∼20-30 minutes, the trial was repeated again for a total of 2 acquisition trials per mouse per day. After all males completed the 2 trials, then female mice were tested with ∼20-30 minutes inter/trial interval. <u>Probe trial:</u> the escape box was removed and the mouse was allowed to explore the maze for 90 seconds, after which the mouse was put back in its individual cage. Only 1 probe trial was performed. <u>Reversal trials</u>: the escape box was moved to the exact opposite location in respect of the target hole location during the acquisition phase. Testing was performed the same way as the acquisition trial but it lasted only 3 days. <u>Reverse probe trial</u>: identical as the probe trials.

### Data analysis

Data was extracted from AnyMaze and analyzed in Microsoft Excel. Plotting and statistical analysis was performed in Prism. Average escape latency was calculated for the 2 trials performed in each day.

### Statistical analysis

Statistical analyses were performed in GraphPad Prism (version 8.4.3) or in R (version 4.2.3). When only two groups were compared, T-test was used if data was normally distributed, or Mann-Whitney if data was not normally distributed, and Welch’s modification was used for data with different standard deviation between groups. When 4 groups were analyzed, we used 2-way ANOVA with either Dunnet or Tukey’s correction for multiple comparison. For sequencing analysis DESeq2 (version 1.38.3) R package was used to determine statistically significant groups. Plotting was done in R using the ggplot2 package or in Prism.

## SUPPLEMENTARY FIGURES

**Supplementary Fig 1 (Related to Fig 1):**
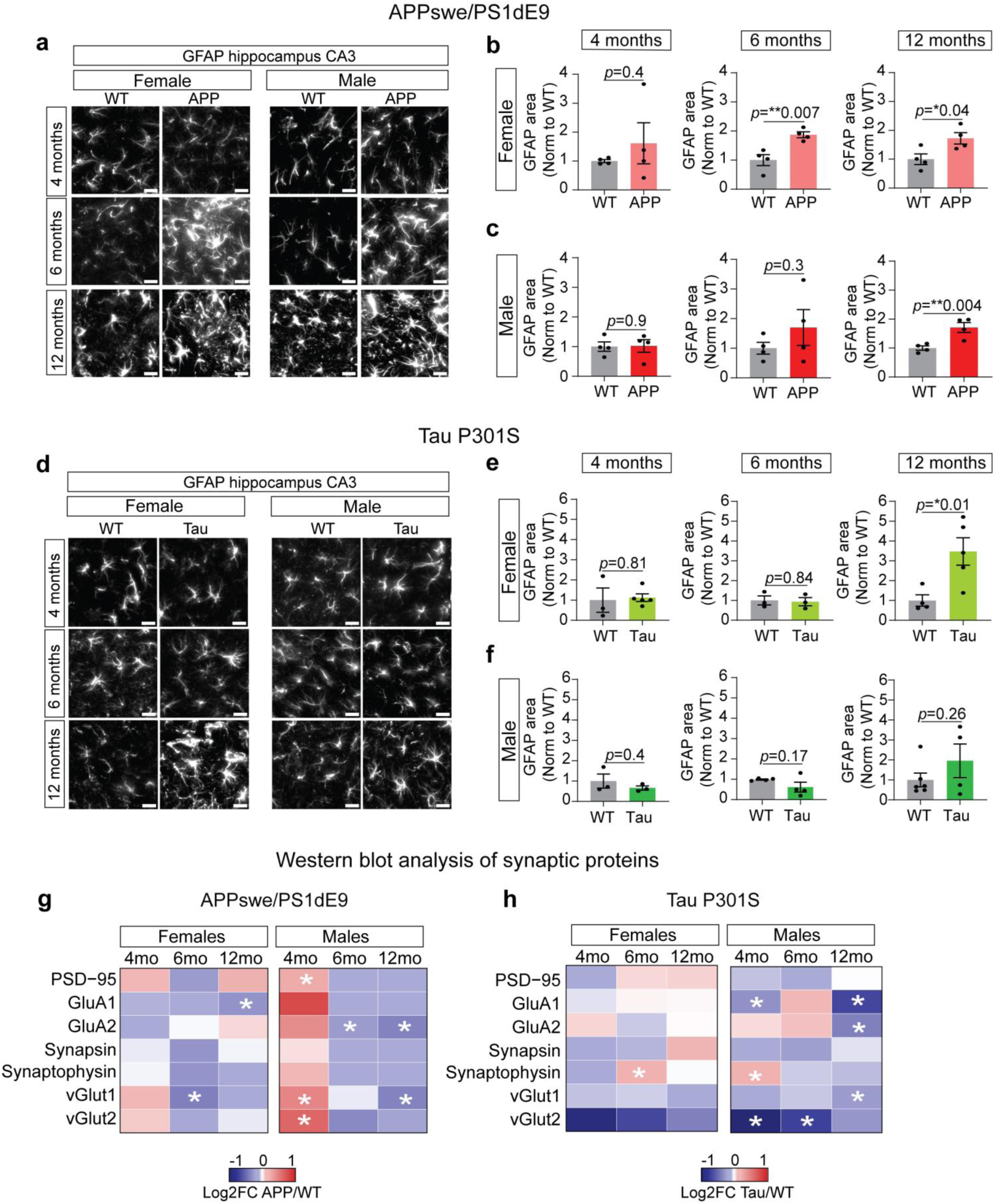
APPswe/PS1dE9 and TauP301S mouse characterization. a-c) Representative images of hippocampal CA3 GFAP immunostainings in male and female mice in APPswe/PS1dE9 mice at 4-6-12 months (a). Quantification of GFAP area in CA3 in females (b) and males (c). Data normalized to WT mice. *N*= 4-5 mice per group/sex/age. Statistical analysis: T-test. Graphs show the mean ± SEM. Scale bar = 20 µm. d-f) Representative images of hippocampal CA3 GFAP immunostainings in male and female mice in TauP301S mice at 4-6-12 months (d). Quantification of GFAP area in CA3 in females (e) and males (f). Data normalized to WT mice. Each data point represents an independent mouse. *N*= 4-5 mice per group/sex/age. Statistical analysis: T-test. Graphs show the mean ± SEM. Scale bar = 20 µm. g-h) Heatmap showing APP (g) and Tau (h) versus WT Log2FC of different synaptic proteins measured by Western blot in the whole hippocampus. *N*= 4-5 mice per group/sex/age. Statistical analysis: T-test, Mann Whitney or Welch test * *p*<0.05.

**Supplementary Fig 2 (Related to Fig 1):**
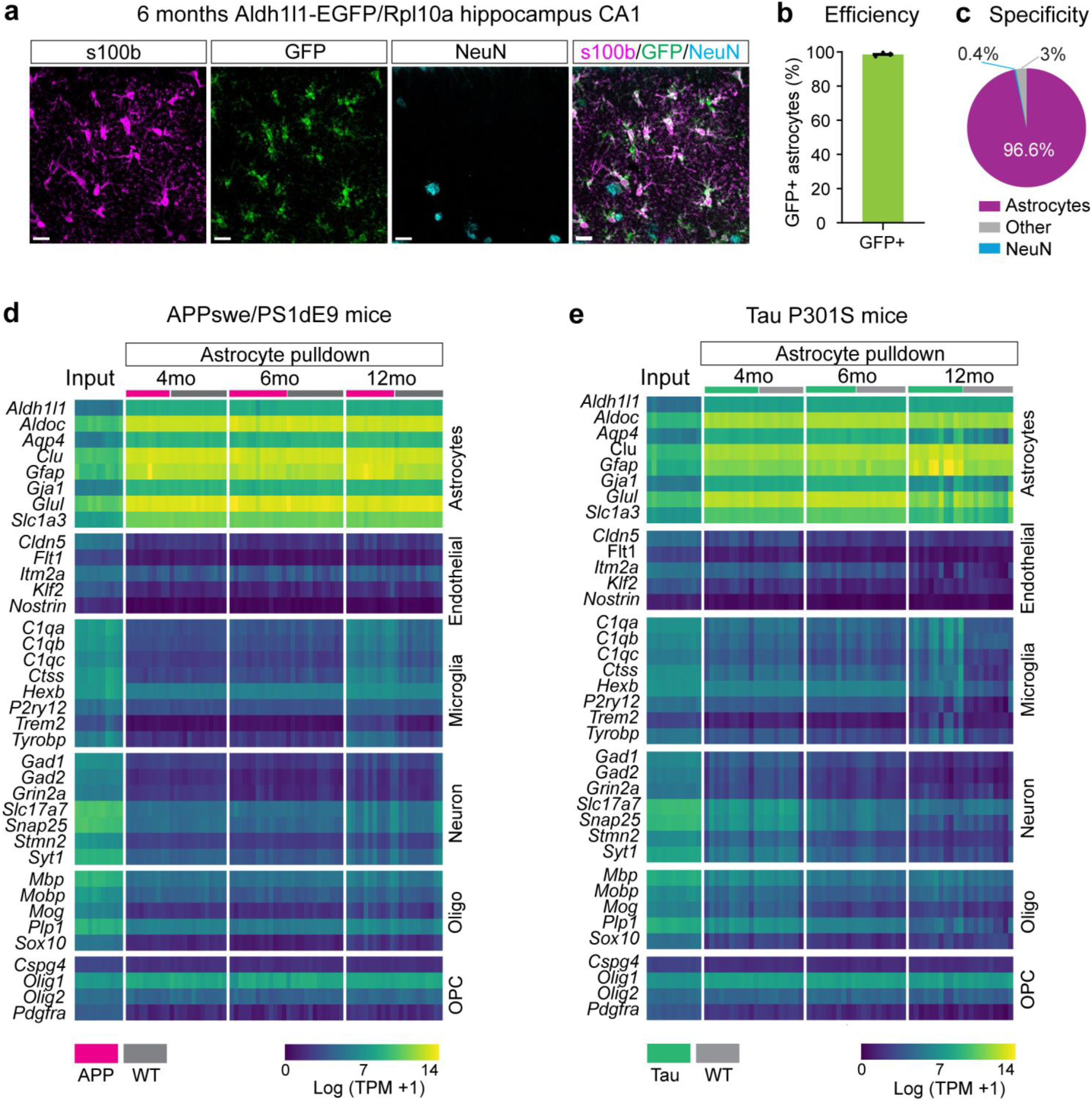
Aldh1l1-EGFP/Rpl10a mouse characterization. a-c) Representative image of 6-month-old Aldh1l1-EGFP/Rpl10a mouse showing GFP colocalization with the astrocyte marker s100b and no colocalization with neuronal marker NeuN. Scale bar = 20 µm (a). b) Quantification showing percentage of s100b+ astrocytes expressing GFP-tagged ribosomes, and c) percentage of GFP positive cells colocalizing with astrocytes, neurons or other cells. d-e) Heatmap showing expression values for different cell type markers in the astrocyte pulldown relative to the hippocampal input for APP (d) and Tau (e) across different ages. Colors represent Log(TPM+1). Olig: oligodendrocytes, OPC: Oligodendrocyte precursor cells.

**Supplementary Fig 3: (related to Fig 4):**
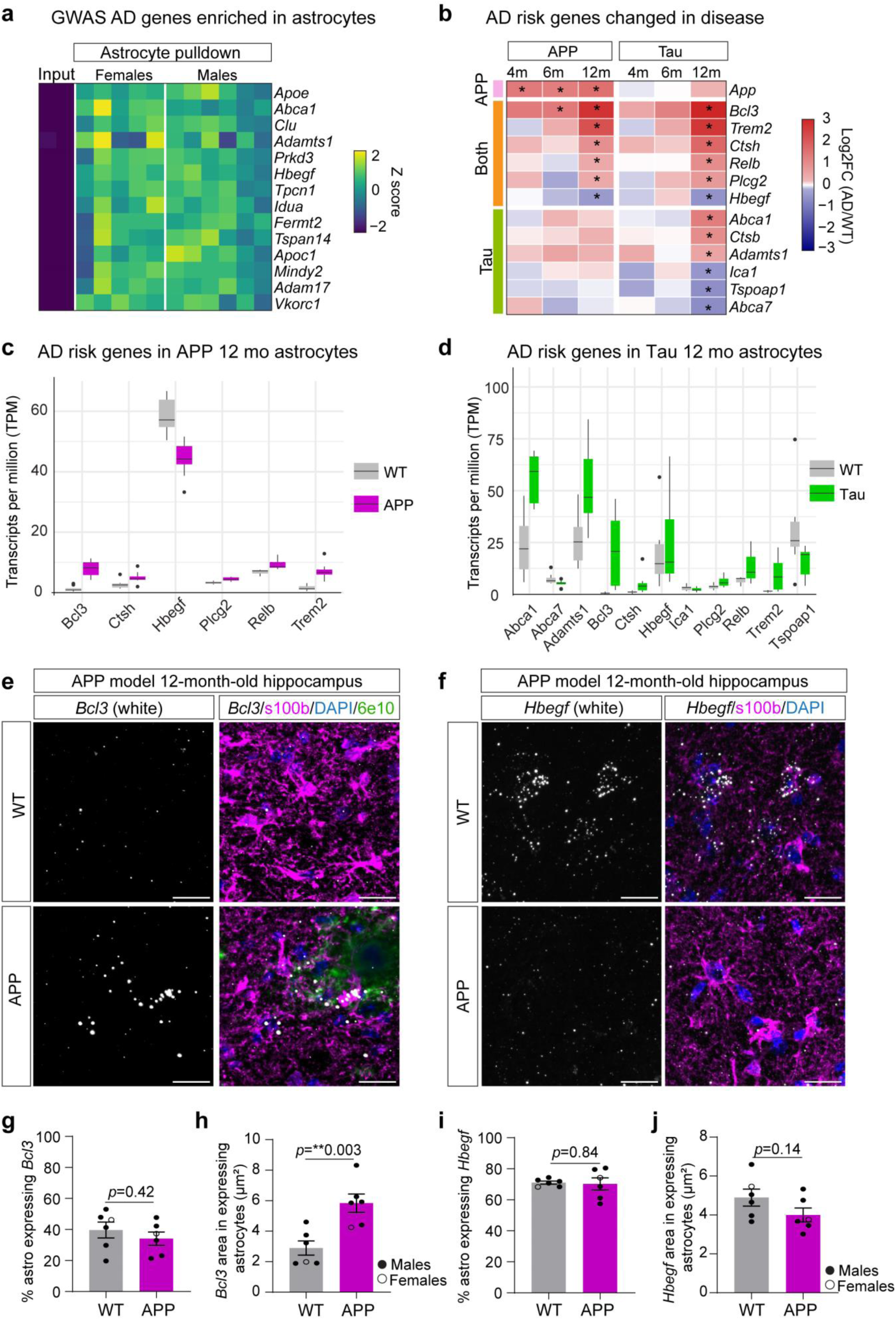
Expression of GWAS AD risk genes. a) GWAS AD risk genes enriched in the astrocyte fraction (Log2FC pulldown/input>1, *padj* <0.05) in 12-months-old WT mice (from the APP dataset). Colors represent expression levels (log (TPM+1) in the different fractions (input versus astrocyte pulldown). *Ace* gene also showed an astrocyte enrichment but was removed due to high variability. Complete list of genes available in Supplementary Table 11. b) GWAS AD risk genes that are significant DEGs in APP and Tau mice at 12 months. Colors represent Log2FC at different ages in the two models. c-d) GWAS AD risk genes that are significant DEGs in APP (c) and Tau (d) mice. Boxplots show the expression levels (TPM) of these DEGs in 12-month-old WT and transgenic mice. Box represents the interquartile range covering the 25th to 75th percentile of the data, with middle line showing the median. Whiskers lines extend to 1.5* interquartile range, and dots represents outliers outside this range. e-f) Representative image of *Bcl3* (e) and *Hbegf* (d) single molecule mRNA *in situ* hybridization co-immunostained with astrocyte marker s100b and amyloid plaques marker 6e10, in 12-month-old WT and APP hippocampus. Scale bar=20 µm. g-j) Quantification of percentage of astrocytes expressing *Bcl3* or *Hbegf* mRNA (g, i) and *Bcl3 or Hbegf* mRNA area in astrocytes that are expressing (h, j). *N*= 5-6 mice, each dot represents a mouse. Graphs show the mean ± SEM. Statistical analysis calculated using T test.

**Supplementary Fig 4: (related to Fig 5):**
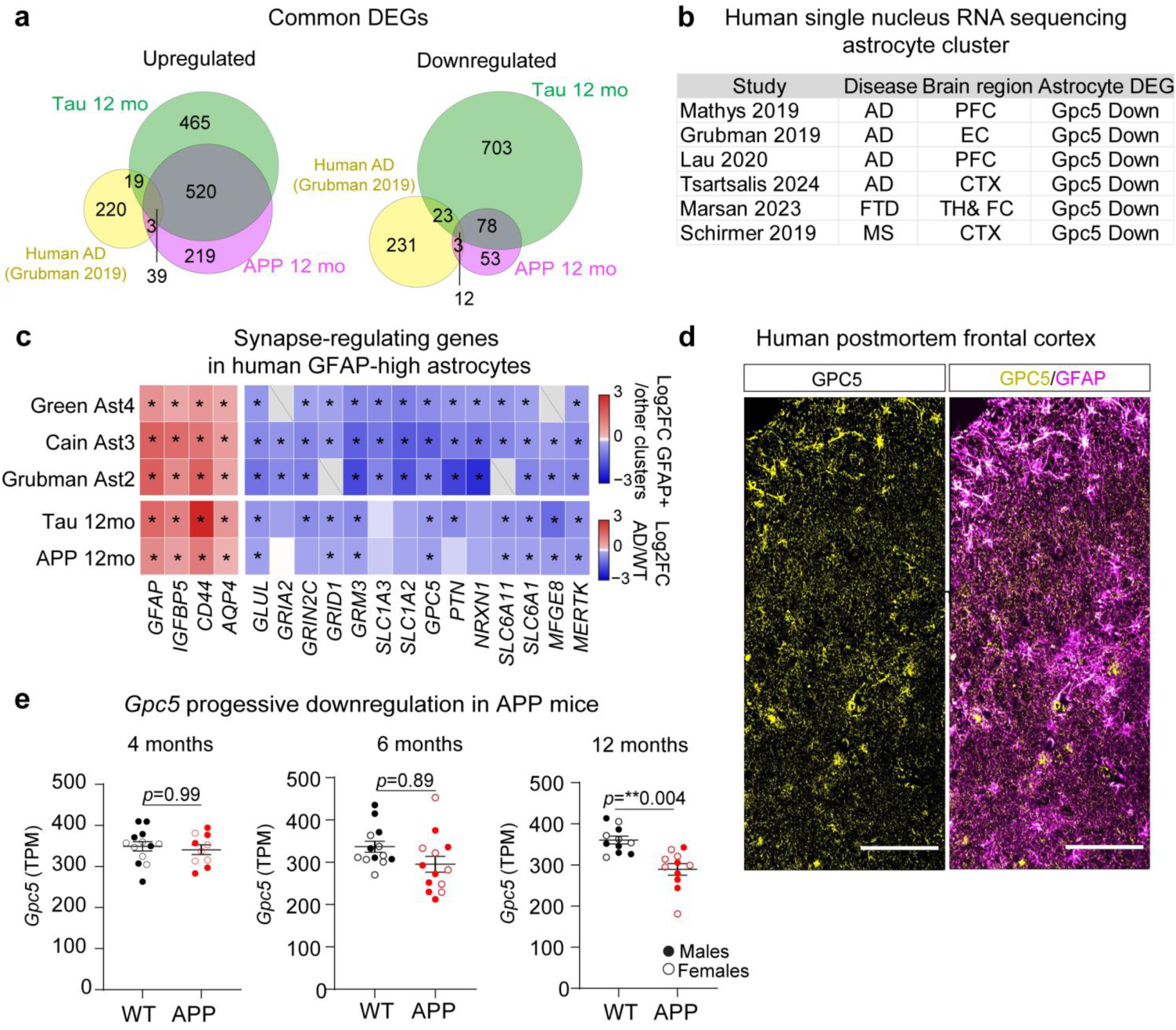
Astrocyte *GPC5* downregulation in AD): a) Venn diagram showing overlapping upregulated (left panel) and downregulated (right panel) DEGs in 12 month-old APP and Tau mice and human AD astrocytes obtained from Grubman et al.^35^ b) Single nuclear RNA sequencing studies from human postmortem patients of different neurodegenerative disorders showing *GPC5* downregulation in the astrocyte cluster. Data obtained from^35–38,43,44^. PFC=prefrontal cortex, EC=entorhinal cortex, CTX=cortex, TH=Thalamus, FC=Frontal cortex. c) Synapse-regulating genes are downregulated in GFAP-high astrocyte clusters relative to other clusters in human AD postmortem brains. Data obtained from^35,40,41^. d) Example image of immunostaining of postmortem human frontal cortex showing GPC5 protein expression in astrocytes marked with GFAP. Scale bar=100 µm. e) Barplot showing TPMs for *Gpc5* expression at 4, 6 and 12 months in APP mice. Statistics show the *padj* value calculated using Deseq2 package in R.

**Supplementary Fig 5 (Related to Fig 6:**
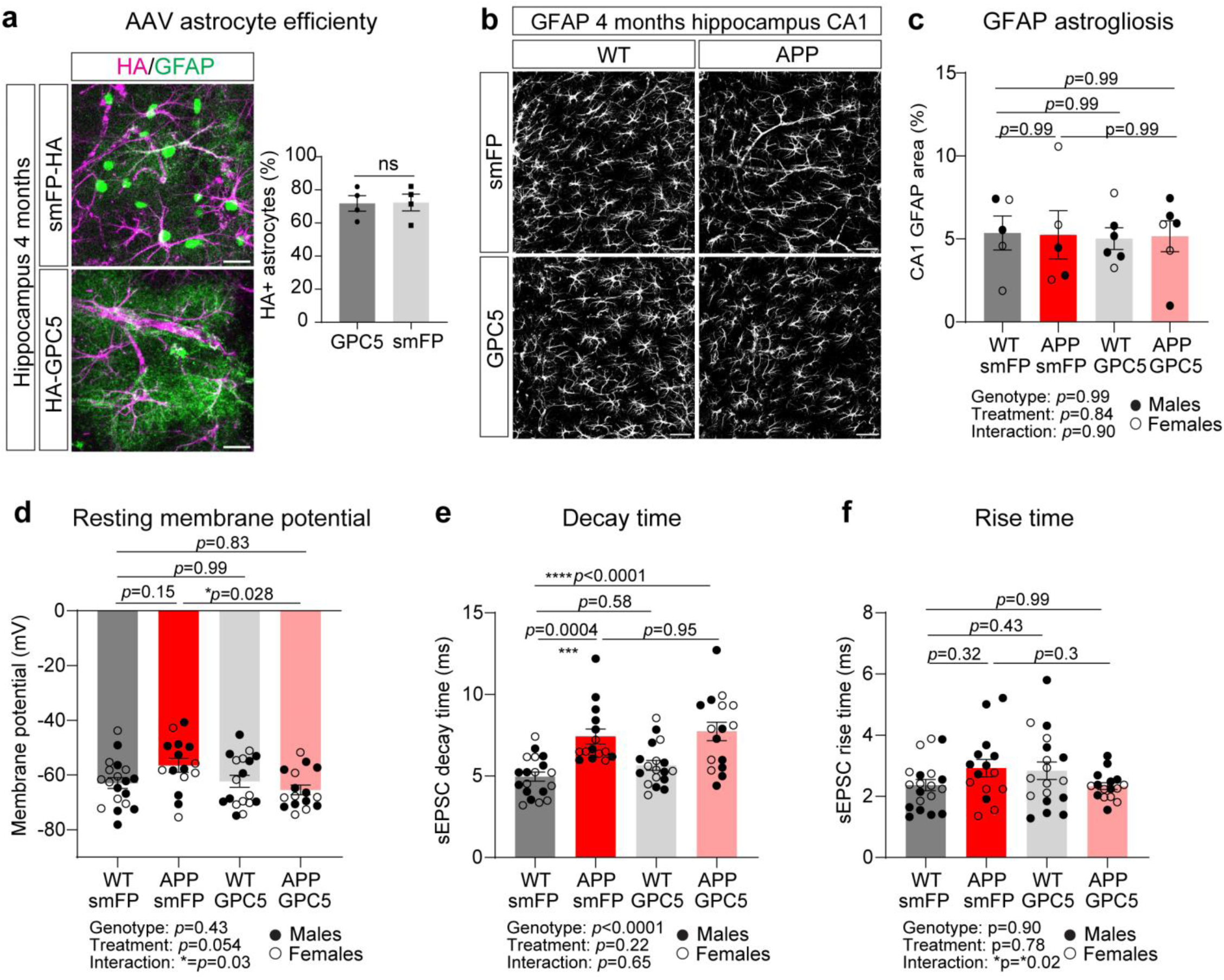
GPC5 overexpression prevents hippocampal synaptic hyperactivity in APP/PS1 mice. a) Representative image (left) and quantification (right) showing colocalization of GFAP-positive astrocytes with HA tag in 4-month-old hippocampus overexpressing smFP-HA or HA-GPC5 for 2 months. Stainings were performed in hippocampal acute slices after electrophysiology recordings were completed. Scale bar=20 µm. Each data point represents an independent mouse. N=3 mice/group. b-c) Representative image (b) and quantification (c) of GFAP staining in the hippocampal CA1 region in WT and APP 4-month-old mice overexpressing HA-GPC5 or smFP control for 2 months. Scale bar=50 µm. Stainings were performed in hippocampal acute slices after electrophysiology recordings were completed. Each data point represents an independent mouse. N=4-6 mice/group. Male (close circles) and female (open circles) mice were included for the analysis. d-f) Electrophysiology recordings of CA1 hippocampal pyramidal neurons performed in 4-month-old APP and WT mice overexpressing HA-GPC5 or smFP showing: d) Resting membrane potential, e) Average decay time and f) average rise time (10-90%) of sEPSC events during 5 minutes recordings. Each data point represents an independent neuron. N= WT smFP: 19 neurons, 9 mice, WT GPC5: 18 neurons, 10 mice, APP smFP: 15 neurons, 10 mice; APP GPC5: 16 neurons, 8 mice. Male (close circles) and female (open circles) mice were included for the analysis. Statistics: 2-way-ANOVA Tukey’s correction for multiple comparison based on neurons. Graphs show the mean ± SEM.

**Supplementary Fig 6 (Related to Fig 6-7):**
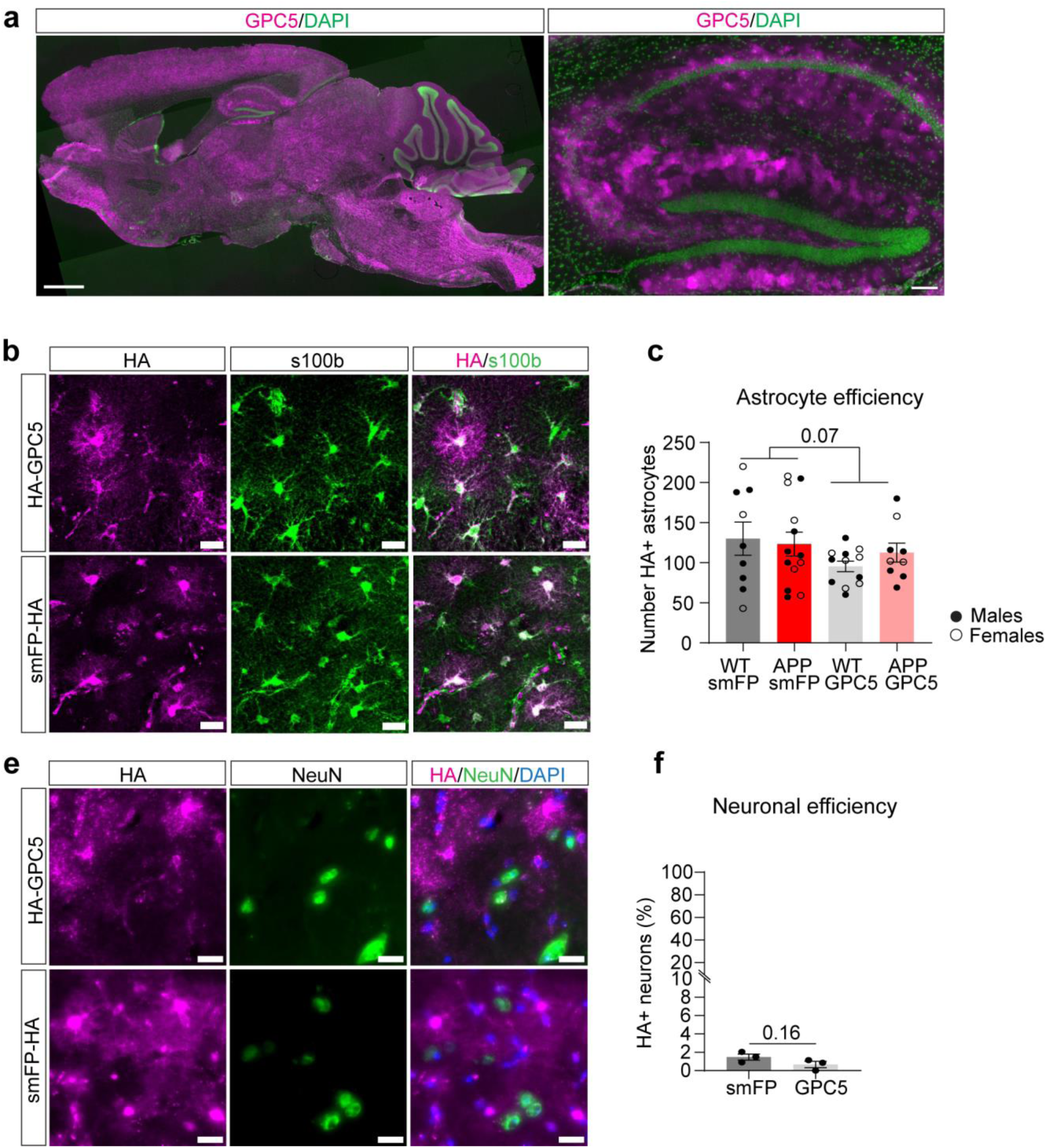
AAV virus characterization. Immunohistochemistry analysis of WT and APP 7-month-old mice overexpressing HA-GPC5 or smFP-HA from 2 to 7 months after testing for memory performance in Barnes maze test. a) Left: Sagittal section of a 7-month-old mouse overexpressing HA-GPC5 for 5 months and immunostained with anti-GPC5 antibody showing GPC5 protein being overexpressed throughout the brain. Scale bar=1mm. Right: Magnification showing hippocampal astrocytes overexpressing GPC5 protein, scale bar=100 µm. Both images were stitched image from a tile scan. b-c) Representative hippocampal image (a) and quantification (b) of 7-month-old mouse overexpressing HA-GPC5 or smFP-HA showing colocalization between HA and the astrocyte marker s100b. Scale bar=20 µm. Same image and quantification were shown in main Figure 6. N= 9-13 mice per group. Male (closed circles) and female mice (open circles) included in the analysis. Statistics: 2-way ANOVA. d-e) Representative hippocampal image showing lack of colocalization between HA and the neuronal marker NeuN in 7-month-old smFP and GPC5-overexpressing mice. e) Quantification showing less than 2% of neurons overexpressing smFP-HA or HA-GPC5. N= 3 mice per group. Statistical analysis: T test.

**Supplementary Fig 7 (Related to Fig 7):**
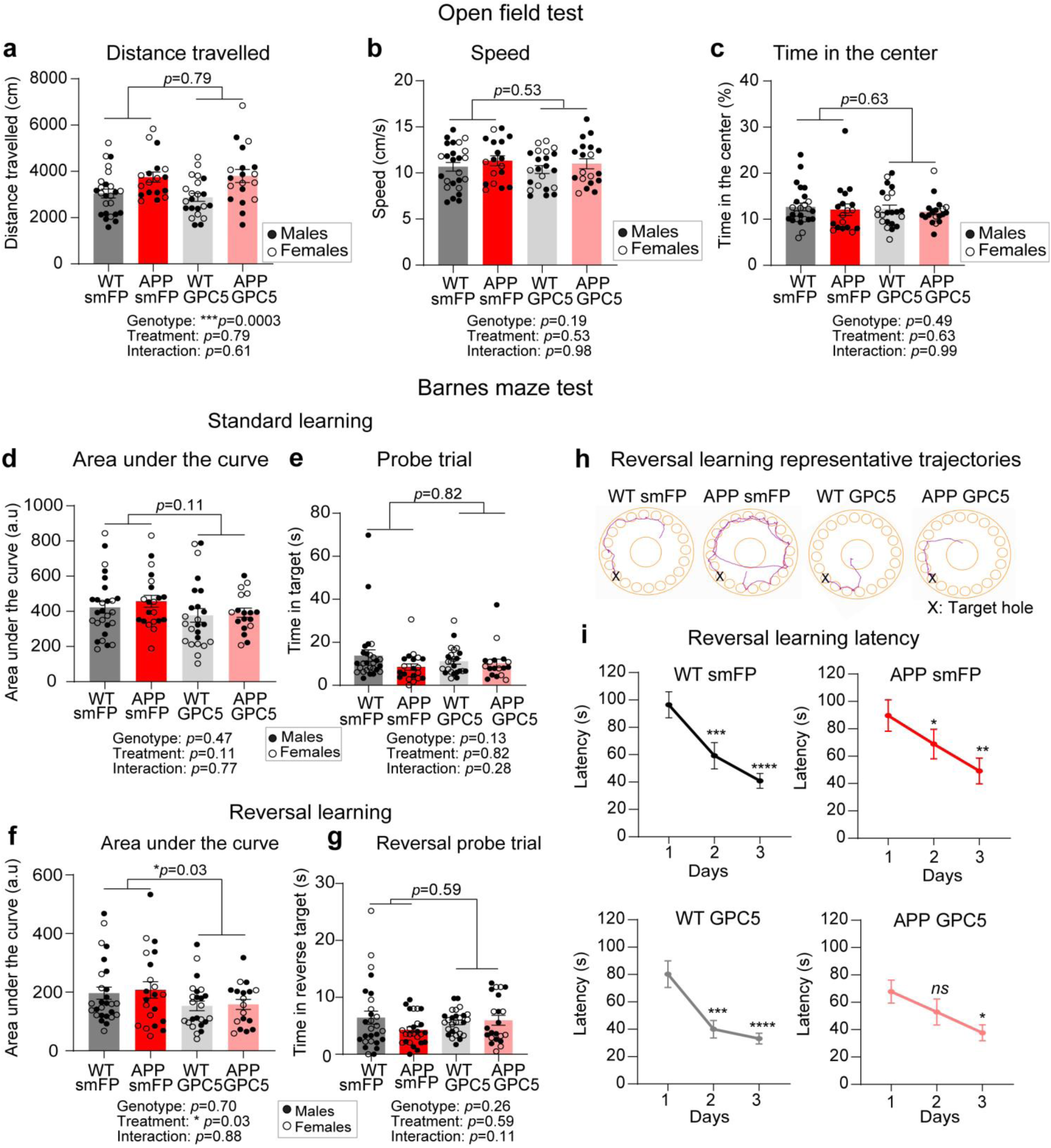
GPC5 overexpression improves reversal spatial learning in APP/PS1 mice. a-c) Open field test consisting of 10 minutes of spontaneous exploration showed an increased total distance travelled in 6-month-old APP mice compared to WT independently of GPC5 overexpression (a) with no changes in mean speed (b) and time spent in the center of the arena (c). d-e) Barnes maze test: area under the curve for the standard learning curve (d) and time spent exploring the target hole during probe trial (e) revealed no differences between groups. Statistical analysis: 2-way-ANOVA. f-g) Area under the curve for the reversal learning curve in the Bares maze test revealed a significant GPC5 treatment effect (2-way-ANOVA, *p*=0.03) (f) and no differences in the time spent exploring the target hole during the reversal probe trial (g). h-i) Representative trajectories used by mice to find the escape box during reversal learning (day 2) (h). Mean latency to find the target hole across reversal learning days (mean of the 2 trials per day). Each panel shows different experimental group. Statistics are calculated using 2-way ANOVA with Dunnett’s multiple comparisons test relative to day 1 (i). Statistics: * p<0.05, ** p<0.01, *** p<0.001, **** p<0.0001. Each data point represents an independent mouse. Male (filled dots) and female (open dots) mice were included for the analysis. WT smFP N=27 (13F, 14M); APP smFP N=21 (10F, 11M); WT GPC5 N=24 (13F, 11M), APP GPC5 N=18 (7F, 11M). Graphs show the mean ± SEM.

**Supplementary Fig 8 (Related to Fig 7):**
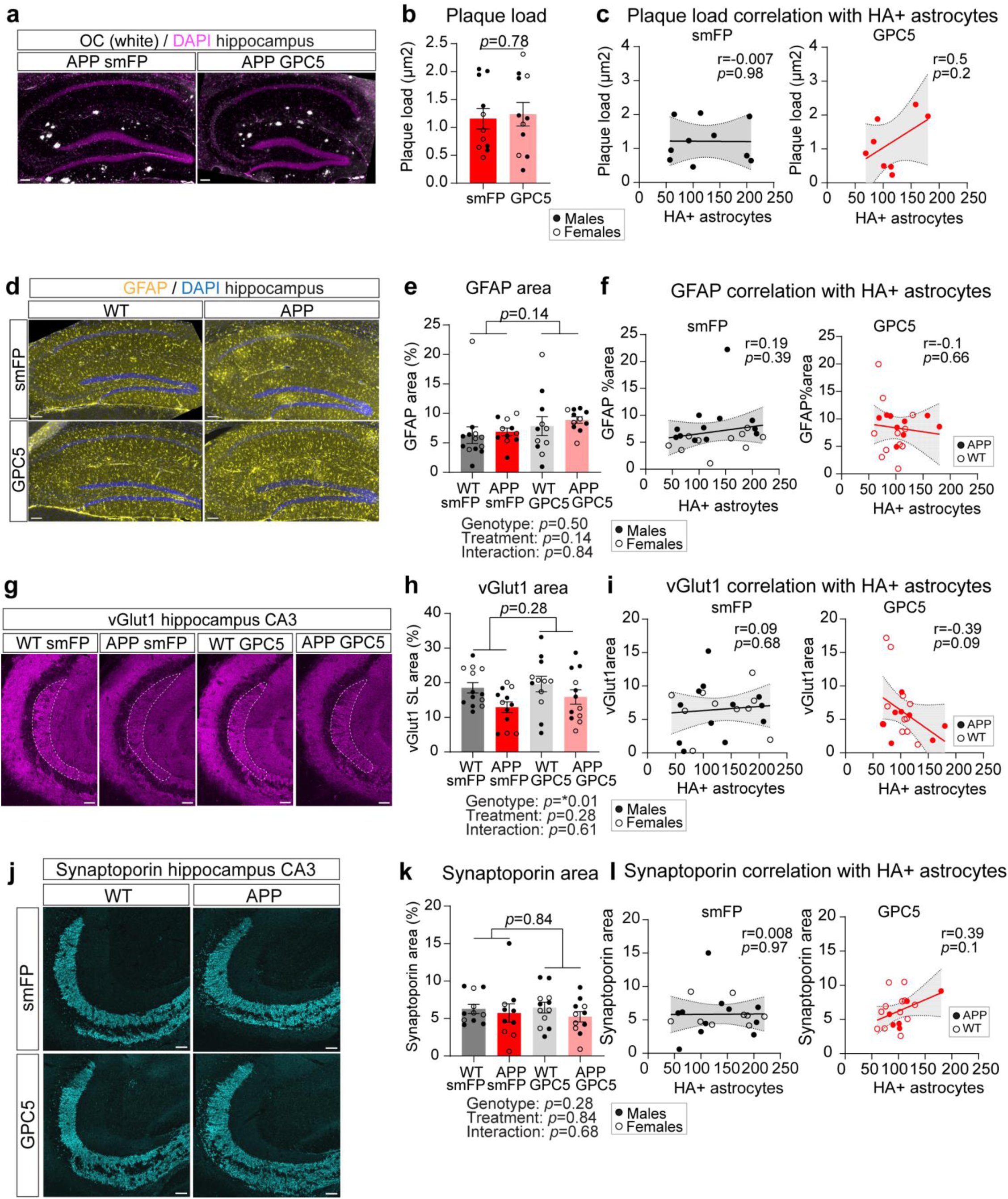
GPC5 overexpression does not affect AD pathology. Immunohistochemistry analysis of WT and APP mice overexpressing HA-GPC5 or smFP-HA from 2 to 7 months after testing for memory performance in Barnes maze test. a-c) Representative tile scans stitched image (a) and quantification (b) of hippocampal amyloid plaques stained with the OC antibody in APP 7-month-old mice. Statistics: T-test. smFP N=11 (5F, 6M); GPC5 N=11 (4F, 7M). Scale bar=100 µm. c) Pearson correlation analysis between plaque load and number of HA-positive astrocytes in smFP-overexpression (left panel) or HA-Gpc5 overexpression (right panel) conditions N=10 smFP, 8 GPC5. d-f) Representative tile scans stitched image (d) and quantification (e) of hippocampal GFAP immunoreactivity in the entire hippocampus across different experimental groups. Pearson correlation analysis between GFAP area and number of HA-positive astrocytes in smFP overexpression (left panel) or HA-Gpc5 overexpression (right panel) conditions (f). WT smFP N=13 (6F, 7M); WT GPC5 N=11 (6F, 5M) APP smFP N=11 (5F, 6M); APP GPC5 N=11 (4F, 7M). Scale bar=100 µm. g-i) Representative tile scans stitched images (g) and quantification (h) of hippocampal CA3 vGlut1 immunoreactivity across different experimental groups. Pearson correlation analysis between vGlut1 area and number of HA-positive astrocytes in smFP-overexpression (left panel) or HA-Gpc5 overexpression (right panel) conditions (i). WT smFP N=13 (6F, 7M); WT GPC5 N=12 (6F, 6M); APP smFP N=12 (5F, 7M); APP GPC5 N=12 (5F, 7M). Scale bar=50 µm. j-l) Representative tile scans stitched image (j) and quantification (k) of hippocampal CA3 synaptoporin immunoreactivity across different experimental groups. Pearson correlation analysis between synaptoporin area and number of HA-positive astrocytes in smFP-overexpression (left panel) or HA-Gpc5 overexpression (right panel) conditions (l). WT smFP N=11 (5F, 6M); WT GPC5 N=12 (6F, 6M); APP smFP N=10 (4F, 6M); APP GPC5 N=11 (5F, 6M) Scale bar=50 µm. Graphs show the mean ± SEM. Each data point represents an independent mouse. Male (filled dots) and female (open dots) mice were included for the analysis. Statistics: 2-way ANOVA Tukey’s test for multiple comparisons, unless specified otherwise.

## SUPPLEMENTARY TABLES

- **Table 1:** Sequencing QC for all groups and exclusion criteria

- **Table 2:** TPM tables 4, 6, 12 months APP and Tau

- **Table 3:** Deseq2 analysis APP

- **Table 4:** Deseq2 analysis Tau

- **Table 5:** Significant DEGs

- **Table 6:** Sex-specific DEGs and pathways

- **Table 7:** Sex-genotype interaction analysis

- **Table 8:** Time-course analysis for DEGs

- **Table 9:** GWAS AD risk genes astrocyte enrichment

- **Table 10:** GSEA functional analysis for APP and Tau

- **Table 11:** Astrocyte synapse-related genes and GSEA

- **Table 12:** Human-mouse overlapping DEGs

